# Incomplete Spatial Patterning of Otic Vesicles in Human Inner Ear Organoids

**DOI:** 10.64898/2026.07.20.739576

**Authors:** Ana Garcia Urbano, Jingyan Yang, David J. Pearton, Mizan Ahmad, Graham Cocks, Alexandre Thiery, Tonglin Zheng, Andrea Streit

## Abstract

During development, patterning of the otic vesicle sets up the blueprint for the adult inner ear. While human inner ear organoids recapitulate the initial steps of ear formation, whether and how vesicles are regionalised remains unknown. Here, we combine re-analysis of published single-cell transcriptomic data with 2D and 3D imaging, using a new PAX2 reporter line, to assess regional patterning of human stem cell-derived otic vesicles. Spanning differentiation days 12-36, we show that some cells co-express markers that are mutually exclusive in vivo, while markers normally occur together are also expressed in different cells. Only a subset of vesicles establishes spatially segregated molecular domains, with patterning poorly coordinated between markers. Adjacent vesicles within the same aggregate differ in their expression patterns, and we found no evidence for a common axis of patterning. These findings point to incomplete resolution of regional programmes and may reflect delayed regionalisation or a lack of the directional and reinforcing signals that pattern the vesicle in vivo.

**One sentence summary:** In human inner ear organoids, otic vesicles show incomplete and disorganised regionalisation, indicating limits of self-organisation under current protocols and pointing to ways of improving regional patterning in vitro.

## Introduction

In humans, hearing loss and balance disorders are among the most common congenital conditions, with lifelong consequences (WorldHealthOrganisation, 2021). Despite this, the mechanisms that govern early human inner ear formation remain poorly understood. During vertebrate development, inner ear progenitors form a simple epithelial vesicle, the otic vesicle (OV) or otocyst, which is progressively transformed into the complex three-dimensional architecture of the adult organ (Alsina et al., 2009; Barald and Kelley, 2004; Groves and Fekete, 2012; Kelley, 2022; Whitfield and Hammond, 2007; Wu and Kelley, 2012). This transformation requires precise coordination of cell fate specification, proliferation, and morphogenesis, and is initiated by OV patterning along the three body axes. Patterning establishes molecularly distinct domains that demarcate future cochlear and vestibular structures, position sensory patches and ensure that cells mature within the correct regional context (Driver and Kelley, 2020; Groves and Fekete, 2012; Kelley, 2022). In vivo, this patterning is directed by coordinated gradients of Retinoic acid (RA), WNT, and SHH emanating from surrounding tissues, which activate distinct transcriptional programmes and ensure reproducible, spatially aligned domain formation (Bok et al., 2005; Bok et al., 2007a; Bok et al., 2007b; Brigande et al., 2000; Chang et al., 2008; Ohta et al., 2010; Ohta and Schoenwolf, 2018; Ohta et al., 2016a, b; Riccomagno et al., 2002; Riccomagno et al., 2005; Whitfield and Hammond, 2007). The mechanisms of OV patterning have largely been investigated in animal models, and it is unknown to what extent these can be transferred to humans.

Over the last decade, inner ear organoids (IEOs) derived from human pluripotent stem cells have emerged as a potential model to investigate such early developmental events in vitro, to model human disease, generate drug-screening platforms and explore hair cell regeneration (Koehler et al., 2017; Liu et al., 2016; Moore et al., 2023; Pianigiani and Roccio, 2024; Tang et al., 2019; Ueda et al., 2023; van der Valk et al., 2021). IEOs generate OV-like structures that give rise to neurons, hair cells, and supporting cells, offering an entry point into human otic development. Recent single-cell transcriptomic studies show that IEOs recapitulate several early features of otic differentiation, including the sequential activation of well-established otic specification genes and cell type specific markers (Doda et al., 2023; Moore et al., 2023; Steinhart et al., 2023; Ueda et al., 2023; van der Valk et al., 2023). However, these analyses also reveal reduced cellular complexity relative to in vivo development and a failure to generate mature cell types, suggesting that organoids do not faithfully recapitulate the entirety of the otic programme (for review: (Nist-Lund et al., 2022; Pianigiani and Roccio, 2024; Rumbo and Alsina, 2024; van der Valk et al., 2021).

In vivo, failure in early ear development leads to defects in its anatomy, cell maturation and function resulting in balance and hearing abnormalities. To increase the fidelity of IEOs in modelling human development it is therefore important to understand when and how they diverge. In IEOs, Wnt pathway activation from differentiation day D8 to D18, first promotes otic placode formation and then generates dorsal, vestibular cell types including immature hair cells, while Wnt inhibition and SHH activation at D12 promotes ventral fates including cochlea-like hair cells (Doda et al., 2023; Moore et al., 2023; Steinhart et al., 2023; van der Valk et al., 2023). Although some regional identity markers have been examined by transcriptional profiling, their spatial distribution over developmental time is not well studied in IEOs. Furthermore, it is unknown whether cells are directly primed towards dorsal or ventral identity, or whether OVs are patterned normally and subsequently lose specific regional characteristics, and, if they are patterned, whether they are organised coherently within each aggregate.

As a first step, we therefore set out to examine axial patterning in dorsal (vestibular) differentiation protocols. We combine the analysis of published single cell transcriptomic datasets (Steinhart et al., 2023) with 2D and 3D imaging using a new PAX2 reporter line to define the extent, variability, and constraints of axial patterning in human inner ear organoids. We show that at the stages analysed (D12-D36) OV-like structures in IEOs can generate regional domains but do so incompletely. While some otic cells express a consistent ‘positional identity programme’, others fail to do so, suggesting that their regional characteristics are not fully established. Finally, we find that there is a strong correlation between patterning and OV size. Together, these findings provide a basis for improving organoid models of early inner ear development.

## Results

### Formation of otic vesicles from human iPCSs

We used a previously reported protocol (Koehler et al., 2017; van der Valk et al., 2026) to generate otic vesicles from the human iPSC line GM25256. As BMP4 levels are critical for non-neural ectoderm induction, we first titrated BMP4 concentrations (0-20 ng/ml, added at D0), scoring aggregate morphology at D3 and the number of OVs aggregate co-expressing otic markers (PAX8/SOX2 and PAX2/JAG1) at D12 (Supplementary Figure 1; 30 aggregates per condition from 3 independent experiments). Epithelial thickness differed significantly between all concentrations tested and 0 or 20 ng/ml BMP4 (Supplementary Figure 1A). Aggregates treated with 1.5-2.25 ng/ml BMP4 had thicknesses within the published range for on-target tissue (38±12 μm) (van der Valk et al., 2026; van der Valk et al., 2023), and 2 and 2.25 ng/ml produced the most OVs per aggregate at D12. We therefore used 2 ng/ml for all subsequent experiments.

Under these conditions, otic differentiation follows similar timelines as previously reported (Koehler et al., 2017; Steinhart et al., 2023). Briefly, at D3, robust expression of the non-neural ectoderm marker TFAP2A was observed (Supplementary Figure 2). FGF treatment together with BMP inhibition at D3 induces pan-placodal fate in the outer layer, with the pan-placodal marker SIX1 being upregulated at D6, together with PAX8, which labels otic-epibranchial progenitors at early developmental stages before becoming restricted to the dorsomedial OV (Doda et al., 2023). Both are strongly co-expressed with TFAP2A at D8 (Supplementary Figure 2). At D6, the neural marker PAX6 is not expressed indicating the absence of any neural tissue (Supplementary Figure 1).

Activation of Wnt signalling by addition of CHIR on D8 marks the start of otic induction, and invaginating placodes can be observed from D9/10 onwards (Figure 1A). This process continues until D18, when CHIR is removed. We visually assessed the emergence of OVs defined by their vesicle-like morphology, and by the expression of at least one otic marker. Around D12, some OVs have already completed invagination and express otic markers. Between D12 and D21, additional vesicles continue to form (see below; Figure 1G). At D18, OVs can still be observed near the aggregate surface, but most have moved towards the interior. Thus, OVs form over a protracted period and aggregates contain vesicles at different stages of maturation.

**Figure 1.**
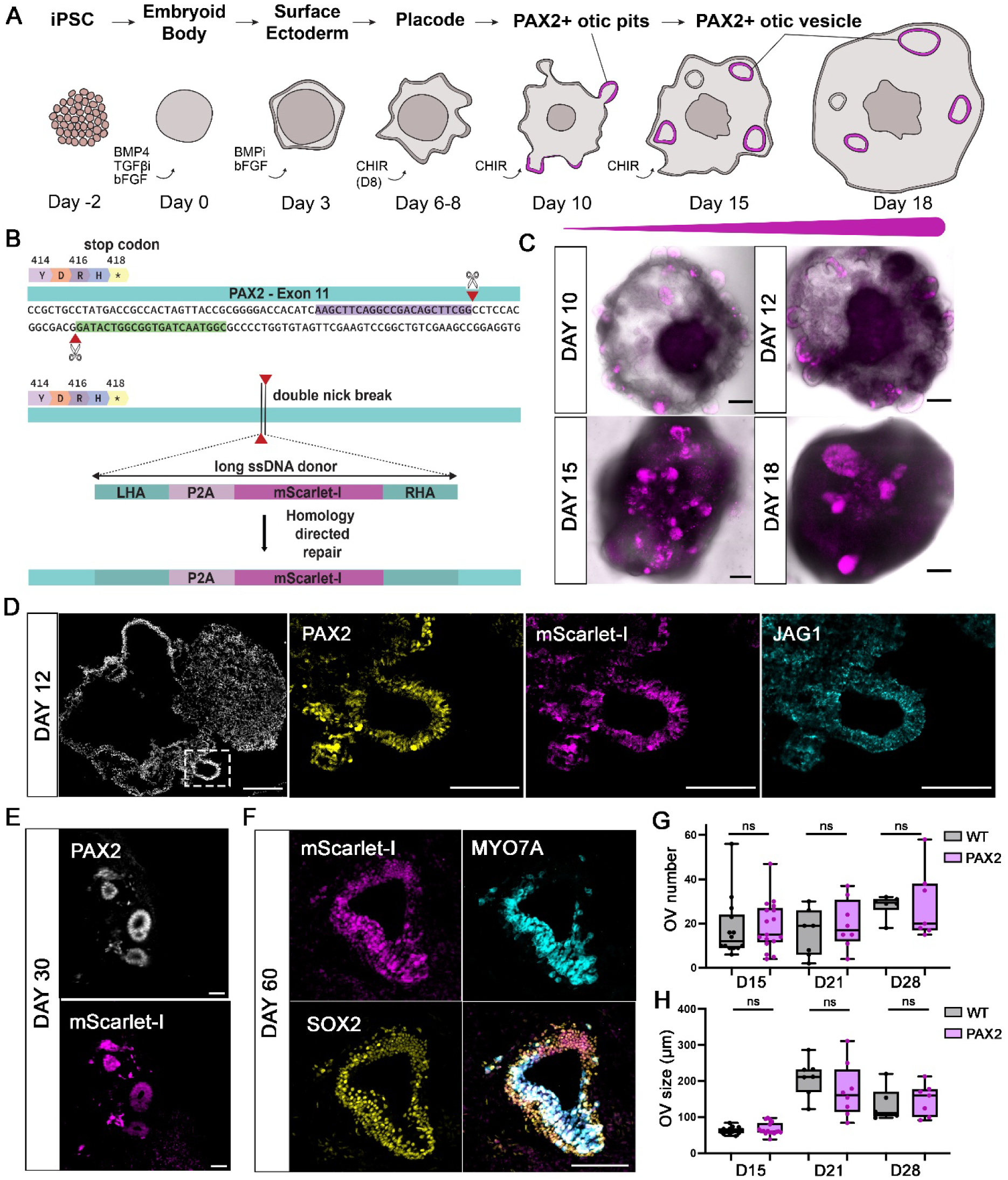
PAX2-P2A-mScarlet-I fluorescent reporter enables tracking of otic vesicles in IEOs. Schematic of inner ear organoid development with PAX2 expression shown in magenta. **B**. CRISPR-Cas9 knock-in strategy: Cas9D10A with two gRNAs (green, purple) introduces paired nicks (red triangles) next to the PAX2 stop codon, and homology-directed repair with a long ssDNA donor (P2A-mScarlet-I flanked by ~1 kb homology arms, LHA and RHA) results in mScarlet-I being transcribed together with PAX2. **C**. Representative live images of aggregates expressing mScarlet-I at days 10, 12, 15, 18. **D**. Expression of PAX2 and JAG1 at day 12. The boxed area in the left image is shown at higher magnification on the right showing an otic pit expressing the pro-sensory marker JAG1 together with PAX2, overlapping with mScarlet-I. **E.** PAX2 and mScarlet-I co-expression in OVs at day 30. **F.** Pro-sensory-like domain in a day 60 organoid containing SOX2^+^ supporting cells and MYO7A^+^ hair cells. Expression of mScarlet-I is maintained in both cell types. **G, H.** Comparison of OV number (G) and size (H) in WT (D15 n=14, D21 n=7, D28 n=6) and PAX2 reporter (D15 n=17, D21 n=8, D28 n=7) lines from 3 independent differentiations, with each IEO representing one biological replicate. Box plots show the median, interquartile range and minimum–maximum whiskers. Statistical comparisons were performed using two-sided Welch’s t-test or Mann–Whitney U test, as appropriate, with Benjamini– Hochberg FDR correction. There is no difference between both lines. Scale bars: 200 μm in C and D (left image), 100 μm in D (higher magnification) and F, 50 μm in E.

At D36, only very few hair cells can be defined transcriptionally (12 of a total of 1,323 otic epithelial cells, 0.9%) (Steinhart et al., 2023). Accordingly, we observe a very small number of ATOH1 positive cells at D36 (not shown) and robust MYO7A at D60 (Figure 1F). Thus, the developmental timeline of our IEO protocol closely matches published data (Doda et al., 2023; Koehler et al., 2017; Steinhart et al., 2023; van der Valk et al., 2026; van der Valk et al., 2023).

### A PAX2 reporter line for tracking otic vesicle formation in human inner ear organoids

The transcription factor PAX2 is one of the earliest markers of otic and epibranchial progenitors (Groves and Bronner-Fraser, 2000; Hidalgo-Sanchez et al., 2000; Streit, 2002; Torres et al., 1996) and remains expressed throughout otic development in chicken and mouse, including during hair cell differentiation (Lawoko-Kerali et al., 2002; Warchol and Richardson, 2009). During OV patterning, *Pax2* expression becomes progressively restricted to ventromedial territories, associated with cochlear outgrowth (Hutson et al., 1999; Lawoko-Kerali et al., 2002; Zou et al., 2006). In human embryos, PAX2 is expressed in the OV from Carnegie stages 11–13 (Doda et al., 2023), and labels otic progenitors in IEOs (Moore et al., 2023). PAX2-reporter lines faithfully recapitulate otic development in IEOs (Doda et al., 2023; Moore et al., 2023).

To enable dynamic and quantitative analyses of OV formation and patterning in human IEOs, we therefore generated a CRISPR/Cas9 knock-in PAX2 reporter line in which mScarlet-I is expressed under the control of endogenous PAX2 regulatory elements (Figure 1B; Supplementary Figure 3). In IEO cultures, reporter activity was first detected after Wnt activation by CHIR99021 on D8, coincident with the emergence of otic placodes and OV-like epithelial structures from D10 onwards (Figure 1A, C). Immunofluorescence analysis revealed a high degree of overlap between reporter signal and endogenous PAX2 protein expression (Figure 1D, E), indicating that the reporter faithfully reflects PAX2 expression.

To assess whether the reporter altered otic differentiation, we compared reporter and parental lines. Temporal expression of key otic markers was similar (Supplementary Figure 2), both lines formed OV-like structures and generated hair cells at later stages (Figure 1D–F), and 3D light-sheet analysis showed no significant differences in OV number per aggregate or OV size at D15, D21 or D28 (Figure 1G, H). These data validate the PAX2 reporter line as a tool for studying otic vesicle formation and patterning in human IEOs.

### Incomplete segregation of regional cell states in IEO-derived otic cells

To investigate whether IEO-derived OVs establish transcriptionally distinct regional identities, we examined published single-cell RNA-sequencing (sc-RNAseq) data generated using the same differentiation protocol (Steinhart et al., 2023). We focused on differentiation days 18–36 (D18-36) corresponding to stages when OV-like structures are present and have begun to show differences in sensory and non-sensory cell states, prior to robust cell differentiation (Steinhart et al., 2023). We extracted cells annotated as ‘otic’ from the original annotation and analysed them using UMAP dimensionality reduction (Figure 2A; Supplementary Figure 4A, B).

**Figure 2.**
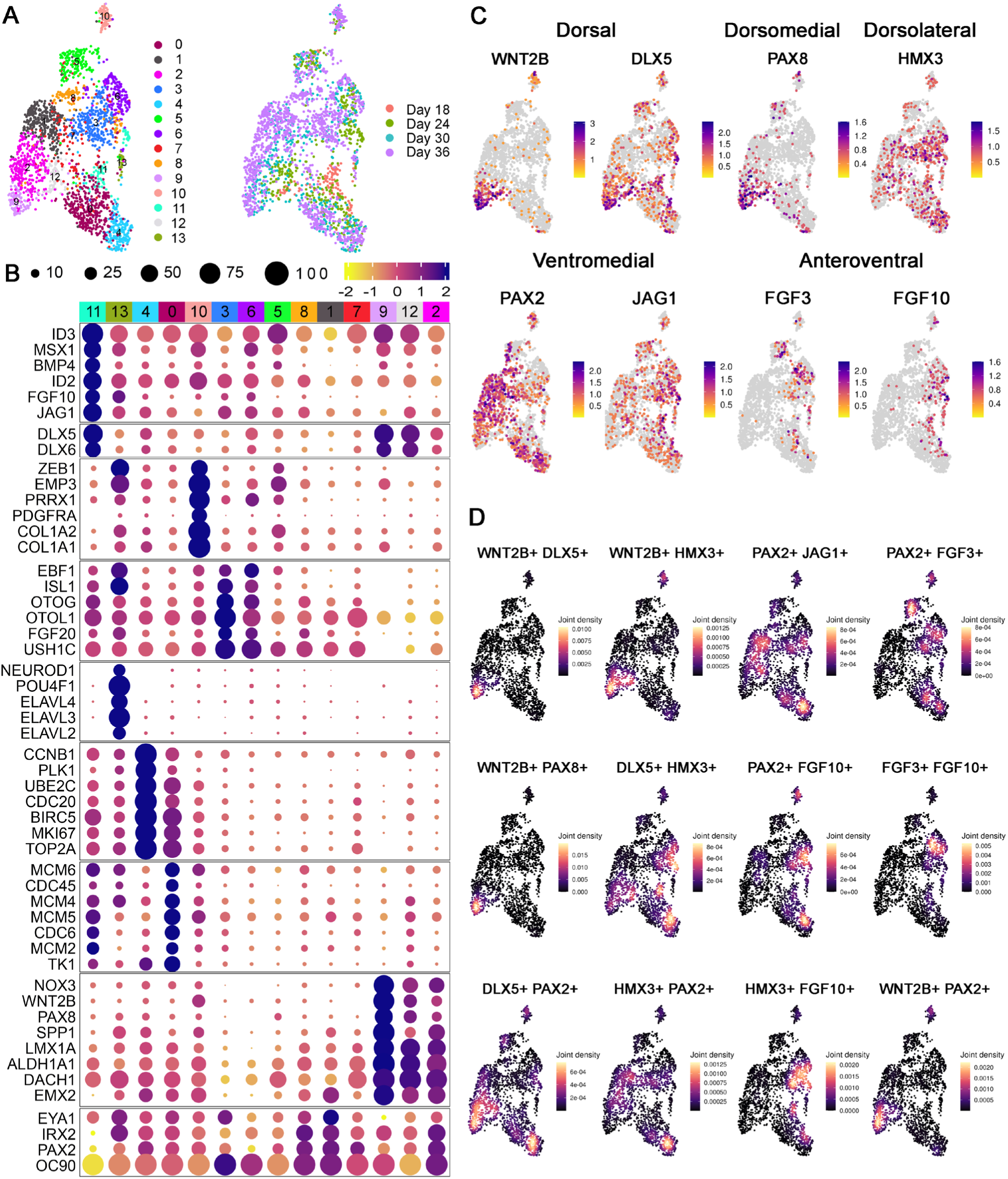
scRNAseq analysis indicates atypical expression of axial markers in IEO-derived otic epithelium. **A.** Otic cells from D18-36 were subset from published scRNAseq data (Steinhart et al., 2023); left: UMAP analysis reveals 14 clusters of cells with similar transcriptional signatures; right: UMAP colour-coded according to differentiation day. **B.** Dotplot displaying the top differentially expressed genes in each cluster. **C**. Feature plots showing the expression of known regional markers in OVs in vivo. **D.** Joint density plots showing pairwise comparisons of the regional markers shown in C. Note: while some regional markers are co-expressed in the same cells as expected, others are not. Likewise, markers showing complementary expression in vivo are expressed in the same cells.

D18-D36 cells segregate into fourteen clusters based on transcriptional similarity (Figure 2A; Supplementary Figure 4). Most clusters contained cells from all differentiation days, except clusters C11 and C12, which are largely composed of D18 and D18-D30 cells, respectively (Supplementary Figure 5). Since only few spatial gene expression patterns are available in human embryos, we compiled a list of markers that define different axial territories of the OV *in vivo* in animal models and human embryos where available (Supplementary Table 1) and assessed whether IEO-derived otic cells segregate based on regional identity. Differential gene expression analysis revealed that clusters C2, C9 and C12 are enriched for genes associated with non-sensory cells (*LMX1A*, *NOX3*, *EMX2*, *DACH1*, *ALDH1A1*; (Steinhart et al., 2023), with C9 showing the strongest enrichment (Figure 2A-C; Supplementary Figure 6; Supplementary Table 2). In addition, C9 and C12 have some dorsal bias with enrichment of *LMX1A, MSX2, DLX5/C* and *WNT2B*. Other clusters are not characterised by regional identity genes (Figure 2A-C; Supplementary Figure 6 Supplementary Table 2). Cluster C3 contains cells with a potential sensory bias (*ISL1*, *EBF1*, *OTOL1*) but cochlear versus vestibular identity cannot be resolved (Figure 2B, Supplementary Table 2). Clusters C0 and C6 represent proliferating cells in S and G2/M phase, respectively, C1 otic-epibranchial progenitor-like cells (*PAX2*, *EYA1*, *OCS0*) and cluster C13 neuroblasts based on the enrichment of neurogenic and neuronal genes (*NEUROD1/4*, *ISL1*, *ELAVL2-4*) (Figure 2B; Supplementary Table 2). C11, the only cluster largely composed of D18-derived cells, is poorly resolved and is split by C13 and contains cells with some neurogenic character (*JAG1, HES5*, *NEUROG1*), while others are characterised by genes related to BMP signalling (*BMP4*, *MSX2*, *ID2/3*). Finally, cluster C10 represents mesenchymal cells potentially of neural crest origin (*COL1A1*, *COL1A2*, *ZEB1 PRRX2*). Four clusters cannot be associated with a clear cell identity. C4 and C8 display a strongly hypoxic profile, C7 has poor marker enrichment with cells being spread across several clusters, while cluster 5 cells are weakly associated with mesenchymal markers (Supplementary Table 2).

To assess the distribution of patterning markers in more detail, we generated feature plots for known axial markers (Supplementary Table 1). The dorsal marker *WNT2B* and the dorsal-medial marker *PAX8* were mostly expressed in C9 (non-sensory, dorsal), while *DLX5* and the dorsolateral marker *HMX3* were detected in multiple clusters (Figure 2C). The ventromedial markers *PAX2* and *JAG1* were broadly distributed, and anteroventral genes like *FGF3* and *FGF10* were expressed in clusters C3 and C6, and C6 and C11, respectively (Figure 2C). Together, these observations suggest that while some otic cells in IEOs are biased towards dorsal identity (C2, C9, C12) they also express genes associated with ventral character suggesting that regional identity is not fully resolved at single cell level (Figure 2C, Supplementary Figure 6).

To better assess gene co-expression in individual cells we generated joint density plots for pairwise comparisons of dorsal and ventral markers (Figure 2D). Cells in cluster C9 indeed co-express the dorsal genes *WNT2B*, *DLX5*, *PAX8* and, to a lesser extent, *HMX3*, but *DLX5* and *HMX3* are distributed widely and co-expressed in individual cells within other clusters (C3, C4, C6, C11). The ventromedial markers *PAX2* and *JAG1* are often co-expressed, overlapping with anteroventral genes like *FGF3* and *FGF10*. However, many individual cells, even in the ‘dorsal’ cluster C9, co-express both dorsal and ventral markers.

Together this analysis reveals that many individual cells co-express conflicting positional identity genes and may therefore remain in an ambivalent transcriptional state. This observation suggests that regional identity of OV cells remains incomplete.

### Spatial expression analysis of patterning genes in IEO-derived otic vesicles

During OV patterning different cell fates are organised in time and space. While scRNA-seq captures the transcriptional states of individual cells, it cannot show whether positional identities are organised into coherent domains. So far, the spatial distribution of patterning markers has not been tested systematically in IEO-derived OVs. We therefore mapped the expression of selected markers in 2D on sections and in 3D by light-sheet microscopy.

#### 2D analysis of regional markers

To provide an initial assessment of the spatial distribution of regional markers, we selected a panel of genes that characterise different OV domains (Figure 3A; Supplementary Table 1) and performed immunofluorescence analysis on cryosections from OV-stage IEOs (D12–36; see Supplementary Table 3 for numbers examined for each marker). As early as day 12, PAX2, SOX2, JAG1 and PAX8 exhibited spatially restricted expression within individual OVs, with patterns generally becoming more pronounced over time. However, not all OVs within the same aggregate shared the same expression patterns, often differing between adjacent vesicles (Figure 3B). Some markers examined exhibited in vivo-like patterns, while others did not. For example, the ventromedial marker PAX2 is often co-expressed with the anteroventral marker SOX2 (Figure 3C; D24) but unexpectedly complementary to the ventromedial marker JAG1 (Figure 3B; D12). PAX2 is also sometimes co-expressed with the dorsolateral marker HMX3 (D21) and the posteroventrolateral marker OTX2 (D30), while we find both complementary and co-expression with the dorsomedial marker PAX8 (D24, D36) (Figure 3B, C). Likewise, SOX2 co-localises with the dorsomedial markers GBX2, PAX8 and LMX1A (Figure 3C D19, D21, D24).

**Figure 3.**
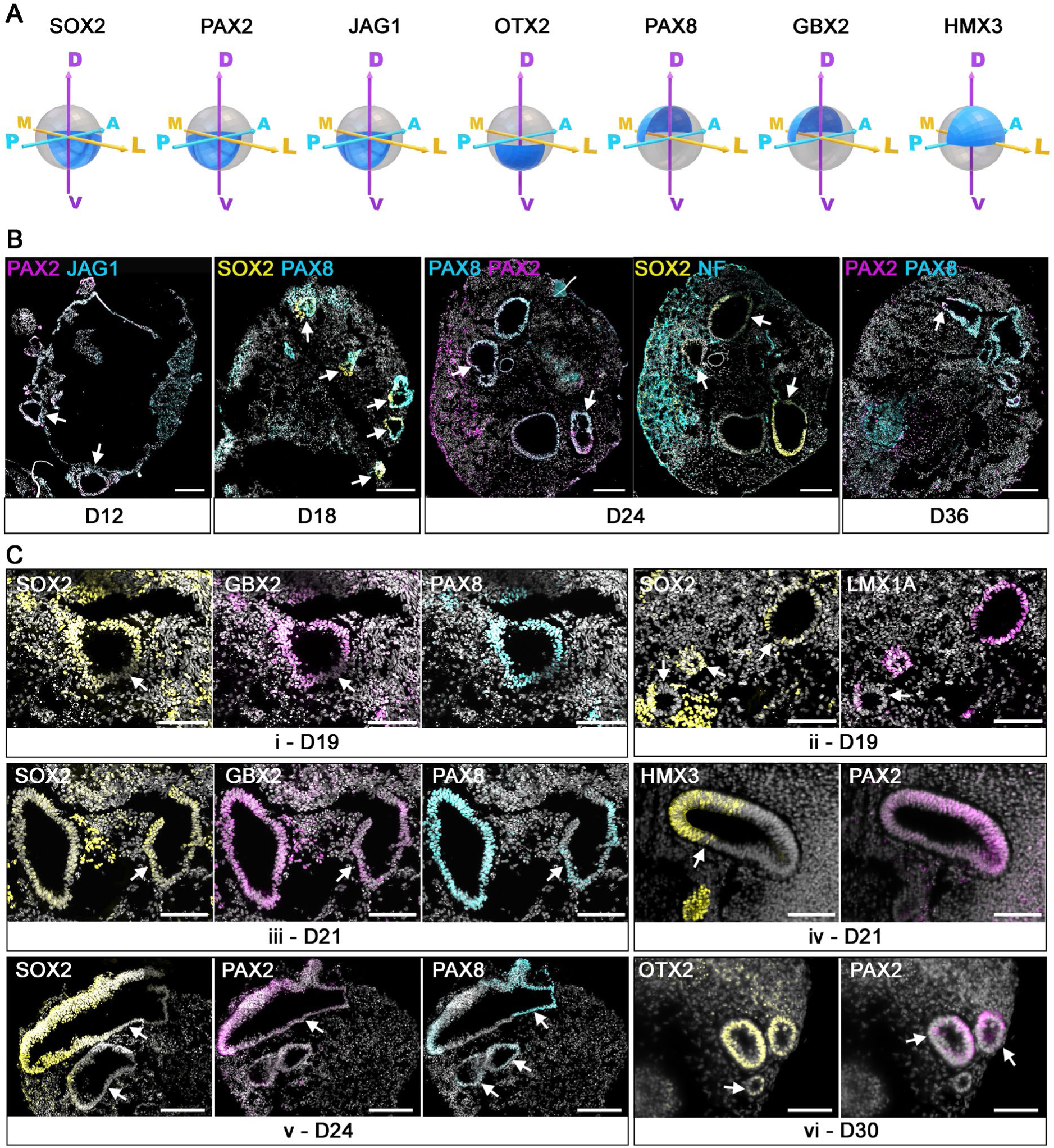
Expression of regional markers in IEO-derived otic vesicles. **A.** Diagrams of otic vesicles showing the expression pattern of regional OV markers as observed in vivo. Note: gene expression is dynamic and markers may change cell type specificity over time (see Supplementary Table 1). The diagrams show the most representative spatial distribution at patterning stages from the literature. **B**. Sections of D12–36 aggregates containing patterned and non-patterned OVs; expression patterns differ between adjacent vesicles. D12: some OVs show patterned JAG1 and PAX2 (arrows), others do not. D18: some OVs show complementary expression of the anteroventral marker SOX2 and dorsomedial marker PAX8 (arrows), but adjacent OVs have different orientations. D24, D36: adjacent OVs show different patterns of PAX8, PAX2 and SOX2. **C**. Higher magnification of individual OVs from D19, D21, D24, and D30. i-D19: overlapping expression of the anteroventral marker SOX2, and dorsomedial markers GBX2 and PAX8. ii-D19: some OVs show complementary expression of SOX2 and the dorsomedial marker LMX1A others show co-expression. iii-D21: adjacent vesicles displaying different expression patterns for SOX2, GBX2 and PAX8; right OV shows a small region of overlap of dorsal and ventral markers (arrow). iv-D21: co-expression of the ventromedial marker PAX2 and the dorsolateral marker HMX3; note: PAX2 is expressed in all cells, while HMX3 is patterned. v-D24: some OVs express SOX2 and PAX2, others do not. The dorsomedial marker PAX8 is expressed in both PAX2/SOX2 positive and negative cells (arrow); note PAX2 and PAX8 staining was carried out on the same section, SOX2 on an adjacent section. vi-D30: co-expression of the posteroventrolateral marker OTX2 and ventromedial PAX2 (arrow). All arrows point to patterned vesicles. Scale bars, 200 μm (B, days 12, 18, 24, 36; C, day 24), 100 μm (C, days 19, 21, 30).

In vivo, OV patterning is initiated by signals from surrounding tissues, and regional identity is later stabilised within the OV itself. To ask whether IEO-derived OVs establish localised signalling domains we examined FGF10 labelling the anteroventral OV and neurogenic domain and WNT2B, a dorsomedial marker (Supplementary Table 1). At D18, OVs did not express FGF10, while surrounding cells with more mesenchymal morphology did (Supplementary Figure 6B). At D21, FGF10 was co-expressed with the anteroventral marker SOX2 in all OV cells. At D18 WNT2B was expressed in all cells at the OV periphery, although the dorsomedial marker PAX8 showed patterned expression, or restricted to a subset of OV cells but co-localising with the anteroventral marker SOX2. Thus, neither marker was localised to the domain expected based on known in vivo patterning.

Overall, these results agree with the notion that the regional identity of individual cells in IEO-derived OVs is not fully resolved and cells continue to express conflicting transcriptional programmes at stages when different OV territories should be spatially defined.

#### 3D analysis of regional markers and quantification of OV patterning

Because sections capture only a single plane of each OV, they provide limited information on OV patterning. We therefore used whole-mount immunostaining and 3D light-sheet microscopy on aggregates collected at differentiation days D15, D21, and D28 (Figure 4; Supplementary Table 5). Within aggregates, individual OVs were defined by their vesicular structure and the expression of at least one otic marker. They were then segmented and analysed for size, marker expression, and patterning status (Figure 4A; Supplementary Table 4). In total we analysed 31 IEOs (570 OVs) from D15, 15 IEOs (268 OVs) from D21 and 13 IEOs (370 OVs) from D28.

**Figure 4.**
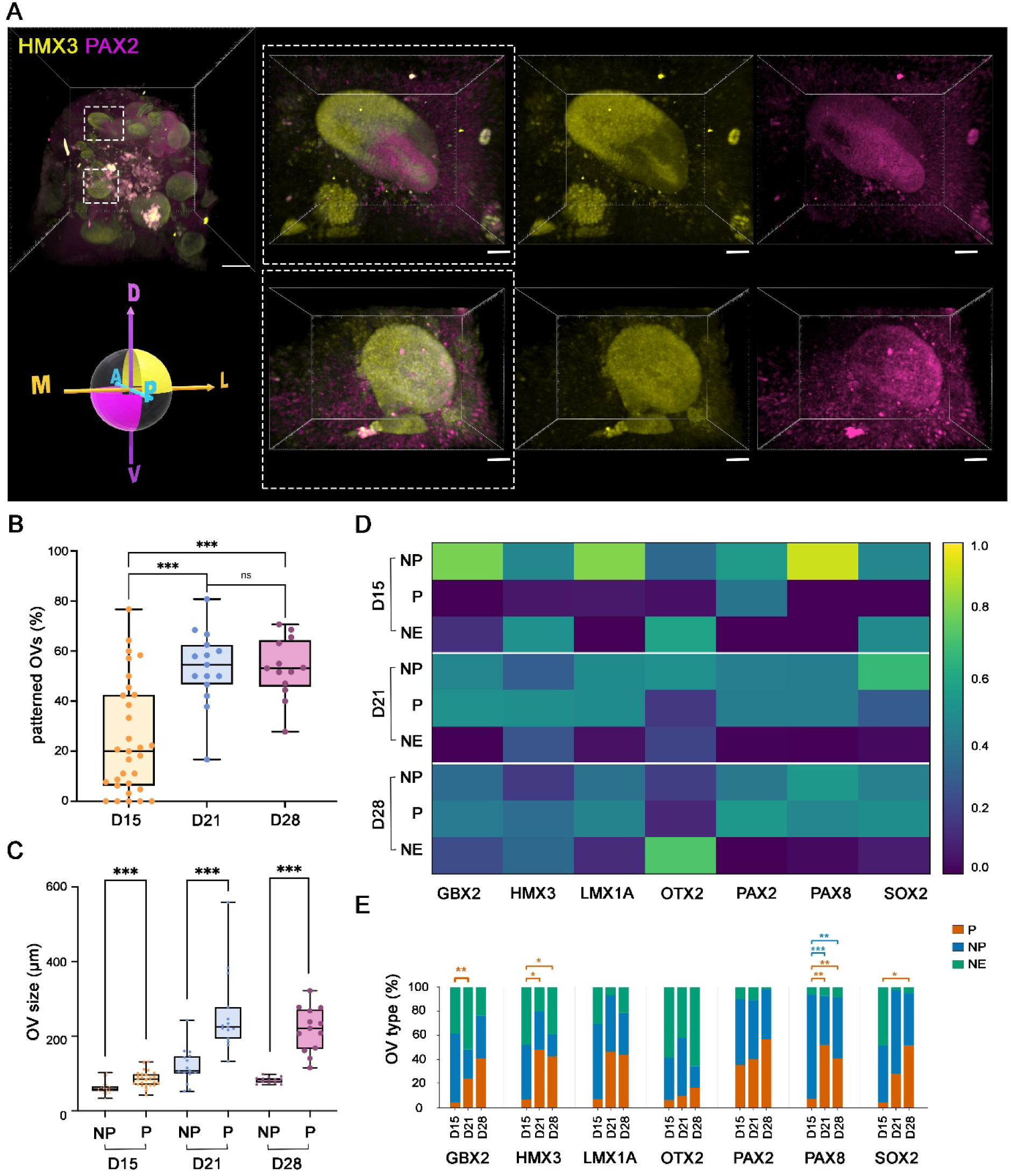
Limited regional organisation in IEO-derived otic vesicles. **A.** D21 aggregate stained for the ventromedial marker PAX2 (magenta) and the dorsolateral marker HMX3 (yellow). OVs within the same aggregate show different expression patterns. Detail of two OVs showing complementary (top) panel and overlapping expression (bottom). **B.** Percentage of patterned OVs per aggregate at D15 (n=31 IEOs), D21 (n=15) and D28 (n=13). Two-sided Mann–Whitney U tests (D15 vs D21, adjusted P = 0.000460; D15 vs D28, adjusted P = 0.000550) and Welch’s t-test (D21 vs D28, adjusted P = 1.000), with Bonferroni correction for three comparisons. ***P < 0.001; ns, not significant. **C.** OV size, defined as the median longest diameter of patterned (P) and non-patterned (NP) OVs within each IEO. Paired, two-sided Wilcoxon signed-rank tests with Bonferroni correction (D15, n = 26 pairs, P = 0.000284; D21, n = 15 pairs, P = 0.000366; D28, n = 13 pairs, P = 0.000732); **** indicates p-value <0.0001. **D.** Quantification of non-expressing (NE) patterned (P) and non-patterned (NP) OVs within each aggregate for each individual marker at D15, D21 and D28. The median for each OV type across all aggregates within the same day and marker category is shown in the heatmap. Values are normalised across groups and shown in a scale 0-1. **E**. Percentage of non-expressing (NE) patterned (P) and non-patterned (NP) OVs for each marker at D15, D21 and D28. Statistical comparisons were performed on IEO-level proportions using Kruskal–Wallis tests followed by Dunn’s multiple comparisons test with Bonferroni correction. Exact adjusted P values are provided in Supplementary Table 4.

While the total number of OVs per IEO increased over time (Supplementary Figure 7A), the proportion of patterned vesicles increased between D15 and D21 but then plateaued at just under 60% (Figure 4B). Likewise, the size of both patterned and unpatterned OVs increased from D15 to D21, and stabilised thereafter (Supplementary Figure 7B, C). Notably, at each stage examined, patterned vesicles were significantly larger than unpatterned vesicles (Figure 4C) suggesting a positive correlation between size and pattern.

Analysing individual markers over time, significantly more OVs expressed PAX2 and PAX8 than any other marker at D15 (more than 90%), which may reflect their early broad expression in vivo before restriction to the ventral and dorsal OV, respectively (Doda et al., 2023; Hutson et al., 1999; Lawoko-Kerali et al., 2002; Torres et al., 1996). Unlike PAX8, PAX2 was patterned in about 30% of OVs compared to under 10% for all other markers, with no significant increase over time. Patterned SOX2 expression occurred in only a few OVs at D15 but increased significantly by D28, whereas OTX2 was expressed in fewer vesicles (less than 50% at any time point) and rarely showed spatial restriction (Figure 4D, E). Most dorsal markers were patterned in less than 10% of OVs at D15, increasing over time; restriction of GBX2, HMX3 and PAX8 increased significantly between D15 and D21 and then plateaued, while LMX1A patterning did not change significantly (Figure 4D, E).

Together, these results reveal considerable heterogeneity between OVs in the same aggregate, with both patterned and unpatterned vesicles present at all stages. Our data suggest that OV patterning correlates with vesicle size, but restricted expression i.e. patterning of any marker does not necessarily correlate with patterning of any other marker. Thus, there is poor coordination of genes that label similar axial OV territories.

### Vector analysis of marker orientation across OVs within individual IEOs

While OV patterning is heterogeneous within IEOs, it is possible that patterned OVs are either oriented along a common axis or preferentially inwards/outwards. To investigate this, we used machine learning assisted 3D segmentation and vector-based analysis (Figure 5A) to analyse the patterning direction of individual markers (PAX2, OTX2, HMX3, PAX8) in one IEO at each of D15, D21 and D28. Because only one IEO was analysed per timepoint, and OVs within an IEO are not independent, we describe the results qualitatively and did not perform statistical tests or compare timepoints.

**Figure 5.**
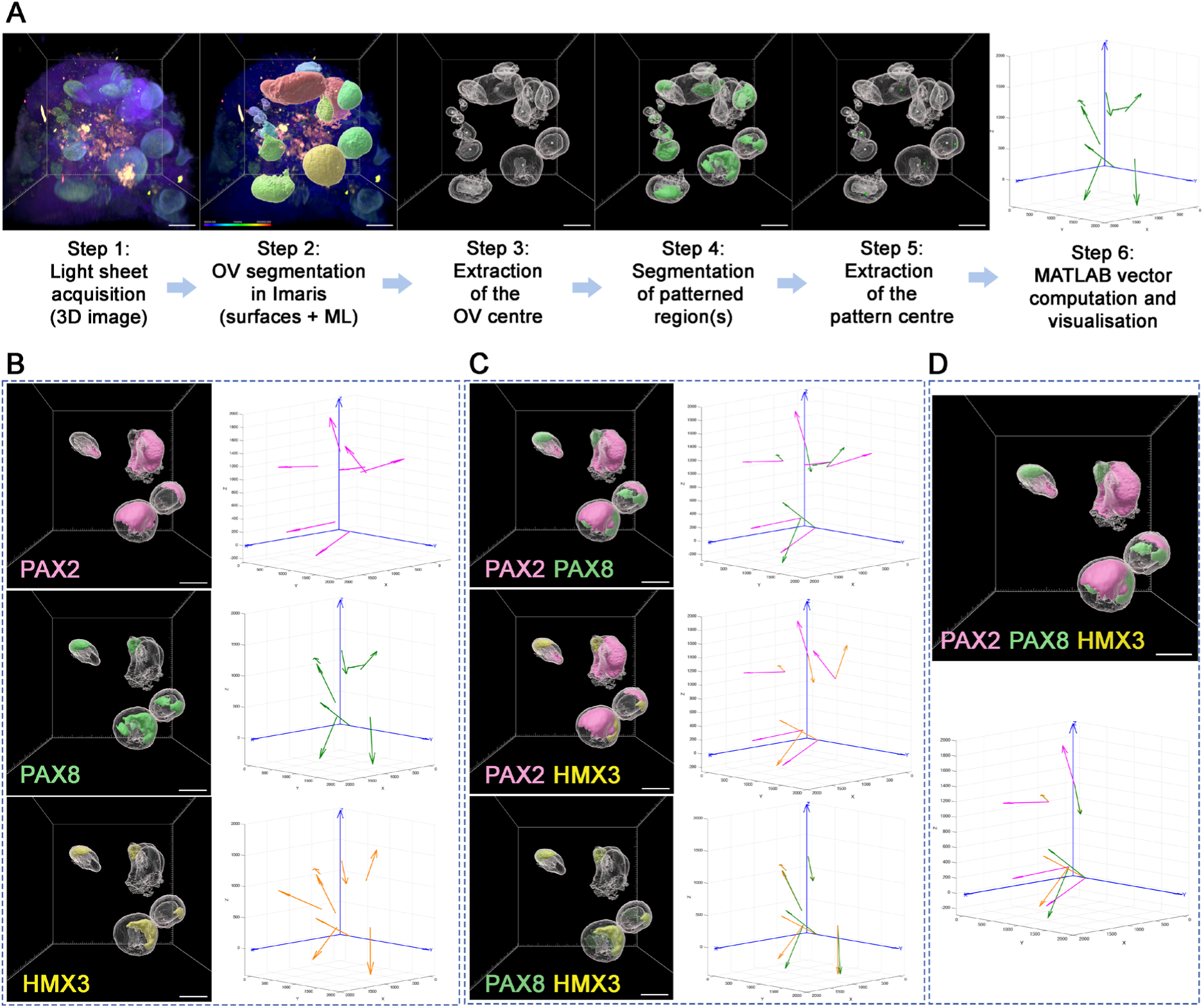
Axial orientation of IEO-derived otic vesicles. This analysis was performed on one IEO for each D15, D12, D28; the figure shows D21 as representative. **A.** Workflow for 3D image analysis of OV patterning. IEO images were acquired by light-sheet microscopy. Individual OVs were rendered as 3D surface in Imaris using a machine learning assisted approach and are shown colour-coded according to surface volume, with colours ranging from purple/blue (smaller) to yellow/red (larger). Marker specific spatial domains were rendered as surface representations. To assess the direction of the patterning axis, we determined XYZ coordinates of the OV and marker domain centres and then used MATLAB to generate vectors linking both. **B**. Representative images showing 3D surface renderings of OVs (grey) with spatial domains of PAX2 (pink), PAX8 (green), or HMX3 (yellow). For each OV within the aggregate, vectors indicate the direction of patterning within each OV. **C**. Pairwise comparison of the direction of patterning between PAX2–PAX8, PAX2–HMX3, or PAX8–HMX3 within the same otic vesicle. **D.** Comparison of three marker domains within the same otic vesicle showing similar direction of PAX8 and HMX3.

To assess overall patterning direction, we represented the orientation of each marker as a vector from the OV centre to the centre of its expression domain (Figure 5A-D; Supplementary Figure 7E-G). We calculated the mean resultant length (R) to summarise the overall alignment of these vectors within each IEO. R close to 1 indicates that vectors point in similar directions, whereas an R close to 0 indicates little overall alignment because vectors pointing in different directions cancel out each other. For comparison, Monte Carlo simulations of randomly oriented 3D vectors gave expected R values of approximately 0.22–0.35 for the numbers of OVs analysed here (7–17). For OTX2, HMX3 and PAX8, R ranged from 0.095 to 0.299, whereas for PAX2 it was 0.133 at D15, 0.483 at D21 and 0.466 at D28 (Supplementary Fig. 7; Supplementary Table 5). All observed R values fell within the central 95% range of the corresponding simulated random distributions.

We also compared the marker-domain vectors of all pairs of OVs within each IEO using cosine similarity (1: same direction; 0: perpendicular; −1: opposite) and calculated the angle θ between vectors (Supplementary Figs. 7, 8; Supplementary Table 5). For descriptive summaries, pairs were classified as similar (θ ≤ 45°), intermediate (45° < θ < 135°) or approximately opposite (θ ≥ 135°); randomly oriented 3D vectors would give about 15%, 71% and 15% of pairs, respectively (see Methods). Across markers and IEOs the proportions were close to these expectations (14-31% similar, 55-68% intermediate, 14-25% opposite; median cosine similarity −0.21 to 0.21). The main exception was PAX2 at D21 and D28, where 29% and 31% of pairs were similar and only 14% were opposite, consistent with its higher R. Thus, these analyses did not indicate a common axis of patterning across OVs, although weak biases cannot be excluded given the small number of OVs.

Because R and the pairwise comparisons assess alignment in a shared coordinate frame, they cannot detect vectors that are consistently directed relative to the position of the OV within the IEO, such as pointing away from the IEO centre. We therefore compared each marker vector with a reference vector from the IEO centre to the OV centre (Supplementary Figure 7D-G); angles below 90° were classified as outward and above 90° as inward (random orientation would give about 50% outward). At D15, nearly all vectors were outward (PAX2 14/15, OTX2 17/17, HMX3 15/15). At D21 and D28, most PAX2 vectors remained outward (6/7 and 10/13), whereas other markers were more variable (D21 PAX8 3/8, HMX3 4/8; D28 OTX2 5/12 outward). Thus, an outward tendency was observed for some marker domains but not consistently across markers and IEOs, and with one IEO per timepoint and 7-17 OVs per marker we cannot determine whether it reflects a general feature of patterning.

Overall, we found no evidence for a common patterning axis and only observed a partial, inconsistent outward tendency, but the small number of OVs and the single IEO per timepoint limit firm conclusions.

## Discussion

During development the inner ear arises from a simple vesicle which is gradually transformed into its adult form. Signals from surrounding tissues subdivide the OV into molecularly distinct territories destined to form different parts of the adult ear (Bok et al., 2005; Bok et al., 2007a; Bok et al., 2007b; Brigande et al., 2000; Chang et al., 2008; Ohta et al., 2010; Ohta and Schoenwolf, 2018; Ohta et al., 2016a, b; Riccomagno et al., 2002; Riccomagno et al., 2005). Any failure of this process leads to ear malformations and consequently hearing loss or vestibular dysfunction (Lin et al., 2005; Merlo et al., 2002; Nakajima, 2015; Ohta and Schoenwolf, 2018; Whitfield and Hammond, 2007). IEOs provide an experimentally accessible system to examine OV organisation and cell differentiation in the context of human ear development (for review: (Nist-Lund et al., 2022; Pianigiani and Roccio, 2024; Rumbo and Alsina, 2024; van der Valk et al., 2021). However, it remains unresolved whether axial patterning, and hence functional regionalisation, is reproducibly established in these 3D systems and, if so, if it is coordinated within each aggregate. We show that only a subset of vesicles generates spatially segregated molecular domains, that individual cells co-express regional markers that are well segregated in vivo, and that there is no evidence for consistent orientation within aggregates. Several explanations could account for the incomplete resolution of regional identity: OVs may not be competent to be patterned at the stages analysed, the cues that set up regional identity in vivo may be absent, misaligned or not reinforced in IEOs, or regionalisation may take longer in human OVs than the time frame analysed here. Irrespective of the cause, patterning appears to be sensitive to stochastic and geometric influences, similar to other organoid systems (Chiaradia et al., 2023; Chiaradia and Lancaster, 2026; Lancaster and Knoblich, 2014a, b; Ohlenschlaeger et al., 2026; Rossi et al., 2018).

### Transcriptional heterogeneity may limit the fidelity of OV patterning and cell maturation

One insight from our transcriptomic analysis is that individual cells within organoid-derived OVs co-express gene combinations that are normally mutually exclusive, while gene pairs that are co-expressed in vivo sometimes fail to be expressed in the same cells. Similar transcriptional “blurring” has been reported in other organoid systems; for example, individual cells in cerebral organoids often co-express transcriptional programmes for different brain regions and are not always spatially segregated (Bhaduri et al., 2020a; Bhaduri et al., 2020b; Kanton et al., 2019; Velasco et al., 2019). In vivo, multipotent progenitors appear to co-express competing transcriptional programmes before they commit to specific fates (Soldatov et al., 2019; Subkhankulova et al., 2023; Thiery et al., 2023), which has been interpreted as ‘transitional states’ in which cells maintain their potential to give rise to different cell types. However, recent spatial transcriptomic analysis reveals that in vivo such transitional states are often found at tissue boundaries, where cells decide between alternative fates (Murtazina et al., 2026; Sampath Kumar et al., 2023). In contrast, our spatial gene expression analysis in IEO-derived OVs often shows large domains of incongruent gene expression rather than cells restricted to boundaries. This disruption of expected transcriptional relationships indicates that lineage-specific regulatory networks are not fully stabilised in IEOs, which may in turn hinder reproducible patterning.

An alternative explanation is that co-expression reflects a failure to downregulate early, broadly expressed genes, or a delay in their resolution, rather than a mixed identity. PAX2 and PAX8, for example, are broadly expressed in otic-epibranchial progenitors and only later become restricted in vivo, and in our IEOs both were detected in more than 90% of OVs at D15, with PAX2 often present throughout vesicles in which HMX3 was patterned. Persistence of earlier transcripts and proteins could therefore produce overlap even once cells have begun to diverge, and human OV regionalisation may take longer than in model organisms. It is also possible that human OVs genuinely have more mixed regional identities than mouse or chick, since knowledge of OV patterning derives almost entirely from animal models. Three observations argue against both a simple delay and a species difference, although they do not exclude them: the proportion of patterned OVs levelled off at just under 60% between D21 and D28, co-expressing cells were not concentrated at domain boundaries, where transitional states are found in other tissues in vivo (Murtazina et al., 2026), and the overlap occurred in large incongruent domains. Failure to downregulate could itself reflect missing repressive or reinforcing signals (see below). Distinguishing between delayed resolution, a species difference and mixed identity cell states will require later timepoints, when differentiation becomes more evident, to test whether overlap resolves, coupled with spatial analysis of human fetal inner ears at comparable stages.

Current IEO differentiation protocols ultimately generate hair cells of either vestibular (dorsal) or cochlear (ventral) character based on the differentiation protocol used. However, they are immature and do not recapitulate the diversity of structure and function observed in vivo (Moore et al., 2023; Steinhart et al., 2023; van der Valk et al., 2023). During in vivo development, hair cell maturation occurs within a defined regional framework (for review: (Driver and Kelley, 2020; Groves and Fekete, 2012). Dorsal vestibular and ventral cochlear territories are induced by WNT/BMP and SHH, respectively (Bok et al., 2005; Bok et al., 2007a; Bok et al., 2007b; Brigande et al., 2000; Chang et al., 2008; Ohta et al., 2010; Ohta and Schoenwolf, 2018; Ohta et al., 2016a, b; Riccomagno et al., 2002; Riccomagno et al., 2005; Whitfield and Hammond, 2007), and within this regional context, hair cells subsequently activate different transcriptional programmes, ion channels and stereocilia architectures. However, mismatch of regional identity, timing and extracellular environment prevents full maturation of hair cells. For example, bypassing normal regional patterning by misexpression of Atoh1 generates new hair cells, but they do not always fully mature morphologically and functionally (Guo et al., 2021; Kelly et al., 2012; Lin et al., 2012; Liu et al., 2012; Staecker et al., 2007; Wang et al., 2024). Thus, hair cell maturation is linked to regional patterning and cellular context. It is therefore possible that failure to resolve OV axial patterning represents a bottleneck that limits terminal maturation of human organoid-derived hair cells.

### Lack of directional and reinforcing cues may lead to disorganised OV patterning

Several factors may contribute to the lack of consistent axial patterning in IEO-derived OVs, including the absence or misalignment of morphogen gradients. In vivo, the OV is embedded in a highly structured embryonic environment that provides directional cues: sonic hedgehog (SHH) from the notochord and floor plate induces ventral OV character, which is counteracted by Wnt and BMP signalling from the dorsal neural tube promoting dorsal identity. The protocol used here generates dorsal, vestibular cell types (Steinhart et al., 2023; van der Valk et al., 2026). However, when exposed to Wnt antagonists and SHH agonists, IEOs express SHH-response genes together with other ventral markers and generate cochlear-like hair cells (Moore et al., 2023). Thus, IEO-derived OVs are competent to respond to signalling cues.

In vivo, extrinsic signals initiate OV patterning, but regional identity and cell fates are subsequently refined and stabilised within the OV itself. This reinforcement may be missing in IEOs: the regional markers FGF10 and WNT2B were not localised to the domains expected from in vivo patterns, which is consistent with, but does not demonstrate, a lack of reinforcement, since we did not measure signalling activity. We also find no evidence for coordinated orientation of patterned OVs across an aggregate, indicating that organoids do not establish a consistent aggregate-level polarity.

Together, these observations suggest that while IEO-derived OVs are competent to respond to patterning cues, the likely cause of OV mis-patterning is a lack of directional signals acting at the appropriate time together with the absence of intrinsic reinforcement, highlighting the need to engineer signalling asymmetries that mimic both temporal and spatial dynamics.

### Optimising IEO protocols

Current IEO protocols recapitulate key steps of ear development and cell type specification, but also generate off-target cells, among them mesenchymal cells potentially of neural crest origin (Doda et al., 2023; Moore et al., 2023; Steinhart et al., 2023; van der Valk et al., 2023). Such cells may provide signals to neighbouring OVs in a spatially unconstrained manner, adding variability to the environment in which patterning takes place. In vivo, neural crest and otic progenitors arise in close association (Streit, 2002) with high levels of Wnt promoting neural crest and low levels enhancing otic specification (Brugmann et al., 2004; Litsiou et al., 2005), canonical Wnt activation, augmented by Notch activity, is subsequently required for placode formation (Jayasena et al., 2008; Ohyama et al., 2006). Fine-tuning Wnt and Notch activity over time, rather than providing a constant level of Wnt agonist, is therefore likely to favour otic over crest identity and reduce off-target cells, creating a more controllable environment in which patterning cues can be applied.

Even with a cleaner starting population, cues applied uniformly to an aggregate are unlikely to be interpreted in the same way by all vesicles. In IEOs OVs form over a protracted period, so any aggregate contains vesicles at different stages of maturation (Moore et al., 2023; Steinhart et al., 2023) (this study). Consistent with this, the proportion of patterned OVs increased between D15 and D21 and then levelled off at just under 60%, and patterned vesicles were larger than unpatterned ones. Synchronising OV formation, combined with delivering cues according to their age, may therefore improve the reproducibility of patterning. The PAX2 reporter line allows individual OVs to be tracked, so the timing of cues could be related to vesicle age. Extending culture beyond the stages analysed here would also show whether patterning is delayed or arrested.

Timing alone is unlikely to be sufficient, because uniform exposure to signalling modulators provides no spatial information. In vivo, the OV is patterned along its dorsoventral axis by Wnt and BMP from the dorsal neural tube and SHH from the notochord and floor plate, the same signals that pattern the neural tube itself. In our IEOs, patterned OVs within the same aggregate showed no consistent orientation, and marker domains did not point consistently towards or away from the aggregate centre, indicating that the aggregate does not impose a common axis on its vesicles. IEO-derived OVs respond to Wnt inhibition and SHH activation (Moore et al., 2023) so delivering these signals asymmetrically is a logical next step. Because the same signals pattern the neural tube, approaches developed for neural tube models, such as microfluidic gradients and synthetic signalling centres or organisers (Luo et al., 2025; Rifes et al., 2020; Xue et al., 2024; Xue et al., 2026) could be adapted to OVs. Whether the larger size of patterned vesicles makes them more responsive to such cues, and whether vesicle size or aggregate geometry can be manipulated to favour this, remains to be determined.

Our analysis focused on dorsoventral and mediolateral markers; anteroposterior patterning was not assessed. In vivo, anteroposterior identity of the OV depends on retinoic acid (RA) signalling (Bok et al., 2007a; Bok et al., 2011), and the current protocol provides no defined source or gradient of RA, although modulating RA signalling influences efficiency and fate decisions in IEOs (Duncan et al., 2025). We cannot exclude that absence of anteroposterior cues contributes to the mixed regional identities we observe, and future work should consider all axes, and the signals that establish them, together.

### Conclusion and future directions

Together, these findings highlight both the strengths and limitations of current IEO models. Some OVs form spatially segregated molecular domains, but individual cells co-express markers that are segregated in vivo, only a subset of vesicles are patterned, and we found no evidence for a common axis within aggregates. Whether this reflects delayed resolution of early gene expression, absent directional cues or lack of intrinsic reinforcement remains to be determined. Controlled delivery of dorsoventral and anteroposterior cues, together with synchronised OV development, are logical next steps, and the measures used here provide baseline readouts for assessing such modifications.

## Materials and Methods

### hiPSC culture

The human GM25256 iPSC line was purchased from Coriell Institute for Medical Research and arrived with a statement of verification and authenticity. This iPSC line was derived from fibroblasts reprogrammed using episomal vectors encoding OCT3/4 with shp53, SOX2, KLF4, LMYC, and LIN28A. For additional validation and testing information refer to: https://catalog.coriell.org/0/PDF/NIGMS/ipsc/GM25256_CofA.pdf Human iPSCs (passage 42-52) were cultured in StemFlex Medium (Thermofisher Scientific, cat. A3349401) on Matrigel hESC qualified (Corning, cat. 354277) coated 6-well plates according to manufacturer’s instructions. At 80% confluency or every 2-3 days, cells were passaged at a split ratio of 1:6-1:10 using Versene (Gibco, cat. 15040066) or 0.5 mM EDTA in DPBS (Sigma Aldrich, cat. D8537). Cells were cultured in sterile conditions in a laminar-flow tissue culture hood using aseptic technique. Cells were regularly tested for mycoplasma contamination. To reduce the risk of spontaneous chromosomal duplications or genetic mutations, passage number was limited to P52.

### Inner Ear Organoid differentiation

Inner ear organoids were generated following established protocols (Koehler et al., 2017; Steinhart et al., 2023) with minor modifications. Briefly, hiPSCs at about 60% confluency were dissociated with StemPro Accutase (Invitrogen, cat. A1110501) and distributed at a final concentration of 25000 cell/ml, onto low-adhesion 96-well U-bottom plates (Thermo Scientific, cat. 174925) (100 μl per well) in StemFlex medium containing 20 μM ROCK inhibitor Y-27632 (Tocris, cat. 1254/10) and Normocin (Invitrogen, cat. Ant-rn2) at a final concentration of 100 μg/ml. After 48-hour incubation (differentiation day 0), aggregates were transferred to a new low-adhesion 96-well U-bottom plate in 100 μl Essential 6 (E6) Medium (ThermoFisher, cat. A1516401) containing 4 ng/ml FGF-2 (Peprotech, cat. 100-18B), 10 μM SB-431542 (Tocris, cat. 1614/10), 2 ng/ml BMP4 (ThermoFisher, cat. PHC9534), and 2% Growth Factor Reduced (GFR) Matrigel (Corning, cat. 354230) to initiate non-neural induction. At differentiation day 3, 25 μl E6 containing 250 ng/ml FGF-2 (50 ng/ml final concentration) and 1 μM LDN-193189 (200 nM final concentration; Stemgent, cat. 04-0074-02) per well was added to the pre-existing 100 μl medium. At day 6, 75 μl fresh E6 medium containing Normocin was added to each well. After an additional 2 days (day 8), 100 μl spent medium was removed and replaced by 100 μl E6 medium containing CHIR99021 (Adooq, cat. A10199-5) at a final concentration of 3 μM to promote otic fate, making a total volume of 200 μl per well. After 48h, 100 μl medium in each well was replaced with fresh E6 medium containing 3 μM CHIR. On differentiation day 12, aggregates were collected and washed with freshly prepared Organoid Maturation Medium (OMM) containing a 50:50 mixture of Advanced DMEM:F12 (Thermo Fisher, cat. 12634010) and Neurobasal Medium (ThermoFisher, cat. 21103049) supplemented with 0.5x N2 Supplement (ThermoFisher, cat. 17502048), 0.5x B27 without Vitamin A (ThermoFisher, cat. 12587-010), 1x GlutaMAX (Gibco), 0.1 mM β-Mercaptoethanol (Gibco) and Normocin at a concentration of 100 μg/ml. Next, the aggregates were plated individually into one well of a 24-well low cell adhesion plate in OMM containing 3 μM CHIR and 1% GFR Matrigel and incubated on an in-incubator orbital shaker at 65 RPM at 37°C with 5.0% CO2. Half medium change was performed at day 15. On day 18, Matrigel and CHIR were diluted by performing a half medium change using OMM without Matrigel and CHIR. From differentiation day 18, half medium changes were performed every three days, including one full medium change once a week. After day 45, half medium changes were performed every two days.

### Quantification of aggregate morphology

Aggregate morphology was assessed at day 3 of differentiation. Circularity, epithelial thickness, and surface ruffling were analysed using ImageJ. Epithelial thickness was measured at three different locations per aggregate, as described in (van der Valk et al., 2023). Circularity was quantified using the built-in ImageJ measurement. Surface ruffling was calculated as the ratio of aggregate perimeter to Feret’s diameter, defined as the longest distance between any two points along the aggregate boundary. This ratio was normalised by dividing by *π* and subtracting 1, such that a perfect circle has a value of 0 and increasing values indicate greater surface irregularity.

### Generation of PAX2-P2A-mScarlet-I reporter line

To track otic progenitors using live imaging, we employed CRISPR-Cas9 technology to generate a PAX2-mScarlet-I reporter cell line in collaboration with the Genome Editing & Embryology Core (GEEC) at King’s College London. gRNAs (5′-CCTATGACCGCCACTAGTTACCG-3′ and 5′ AAGCTTCAGGCCGACAGCTTCGG-3′) targeting the stop codon region were designed in silico using the Sanger CRISPR finder tool and the Integrated DNA Technologies (IDT) tool. An mScarlet-I intermediate construct (P2A + mScarlet-I) was kindly provided by Dr Graham Cocks. To generate the PAX2-P2A-mScarlet-I insertion cassette, two ~1 kb homology arms flanking the PAX2 stop codon were PCR-amplified from GM25256 genomic DNA. Gibson assembly was used to join the two homology arms with P2A-mScarlet-I sequence, generating the final insertion cassette. To produce ssDNA donors, we used the Guide-it Long ssDNA Production System v2 (Takara Bio, Cat. 632666). Template dsDNA was generated by PCR amplification of the insertion cassette. gRNAs and Cas9D10A RNP complexes were prepared according to manufacturer’s instructions and transfected together with the ssDNA donor into GM25256 iPSCs using a 4D-Nucleofector with the P3 Primary Cell 4D-Nucleofector X kit (Lonza Bioscience, cat. V4XP-3032) and Program CA-137. Following nucleofection, cells were incubated in StemFlex media containing Revitacell (1:100) to improve cell survival and allowed to recover. Clonal cell lines were then established by low-density seeding (1 – 2 cells cm^-2^) of dissociated single iPSCs followed by isolation of iPSC colonies after 15-20 days of expansion. Genomic DNA was extracted from individual clones, and genotypes were analysed by PCR amplification followed by gel electrophoresis and Sanger sequencing. We obtained one clone with correct P2A-mScarlet-I integration, which was karyotyped and used for subsequent experiments.

### scRNA-Seq reanalysis

The processed Seurat scRNA-seq object described in Steinhart et al., 2023 was downloaded from GEO (GSE188936) as an R object (RDS) and analysed in Seurat V5 (Hao et al., 2024). The preexisting annotations for the full dataset were used. Cells were further subset based on identity, i.e. cells classified as otic by Steinhart, et al., and time, cells from day 18-36 were kept, corresponding to stages when OV-like structures are present.

The D18-36 otic cells were normalised using SCTransform and the Uniform Manifold Approximation and Projection (UMAP) was used for dimensionality reduction. Optimal cluster resolution was determined by clustree analysis (Zappia and Oshlack, 2018) of Leiden clustering with resolutions from 0.4 to 2.0 (Supplementary Figure 4). Differentially expressed genes for each cluster were identified using the Wilcoxon Rank Sum test in FindAllMarkers in Seurat V5 and filtered based on a percentage difference in expression of at least 60% between groups. Featureplots, dotplots and UMAP projects were produced using Seurat V5, scCustomise (Marsh, 2021) and ggplot2. Cluster identity was assigned based on top differentially expressed genes (Supplementary Table 2) and known markers.

### Immunohistochemistry

On collection days, aggregates were washed 3 times with phosphate buffered saline (PBS) and fixed in 4% paraformaldehyde (PFA) in PBS for 20 minutes at room temperature. The fixed aggregates were washed with PBS and cryoprotected with a graded treatment of 15% and 30% sucrose and then embedded in OCT embedding matrix (Fisher Scientific) and frozen at −80°C overnight. Frozen tissue blocks were then cryo-sectioned at 14μm on a Bright OTF5000 cryostat and mounted in SuperFrost Plus™ Adhesion slides (ThermoFisher).

For immunostaining, sections were washed in PBS to remove OCT, incubated in blocking buffer (10% goat serum and 0.1% Triton X-100 in PBS) for 1 hour at room temperature and incubated overnight at 4°C with primary antibodies diluted at the appropriate concentration in PBS containing 5% goat serum and 0.1% Triton X-100. Sections were then washed 3 times with PBS containing 0.1% Triton X-100 (PBST) and incubated with secondary antibodies diluted in 1:500 in PBS containing 5% goat serum and 0.1% Triton X-100. Alexa Fluor conjugated anti-mouse, rabbit, or goat IgG (Invitrogen) were used as secondary antibodies. Cell nuclei were stained with DAPI (1μg/ml) (Invitrogen, cat. D35710). Finally, sections were washed 3 times in PBS and mounted using Fluoromount-G (Invitrogen, cat. 00-4958-02) and glass coverslips. Sections were imaged using a Zeiss LSM980. Images were processed using Zeiss Zen Blue or ImageJ software.

| Antibody | Host | Supplier | Catalogue No. | Dilution |
|---|---|---|---|---|
| OCT 4 | Rabbit | Abcam | Ab181557 | 1:100 |
| SOX 2 | Mouse<br>Rabbit | BD Biosciences<br>Abcam | 561469<br>Ab97959 | 1:100<br>1:100 |
| E-cadherin | Mouse | BD Biosciences | 610181 | 1:250 |
| N-cadherin | Mouse | BD Biosciences | 610920 | 1:100 |
| TFAP2A | Mouse | DSHB | 3B5 | 1:5 |
| PAX6 | Mouse | DSHB | PAX6 (RRID AB_528427) | 1:5 |
| SIX1 | Rabbit | Sigma-Aldrich | HPA001893 | 1:400 |
| PAX8 | Rabbit<br>Mouse | Abcam<br>ThermoFisher | Ab97477<br>MA1-117 (clone IF8-3A8) | 1:100<br>1:100 |
| PAX2 | Rabbit | Zymed | 71-6000 | 1:100 |
| Jagged-1 | Mouse<br>Rabbit | Santa Cruz | Sc-390177<br>2620 (clone 28H8) | 1:50<br>1:100 |
| MYO7A | Rabbit | Proteus | 25-6790 | 1:100 |
| Neurofilament | Mouse | Abcam | Ab64300 | 1:100 |
| m-Scarlet | Goat | St John’s Laboratory | STJ140286-150 | 1:100 |
| LMX1A | Rabbit | Abcam | Ab138726 | 1:50 |
| HMX3 | Mouse | DSHB | PCRP-HMX3-4E5 (RRID AB_2618712) | 1:25 |
| OTX2 | Rabbit | Abcam | Ab9566-1 | 1:250 |
| GBX2 | Goat | Novus Biologicals | NB-100-55393 | 1:100 |
| SOX10 | Mouse | Santa Cruz | Sc-365692 | 1:50 |
| FGF10 | Rabbit | Merck Millipore | ABN44 | 1:100 |
| WNT2B | Rabbit | Atlas Antibodies | HPA066368 | 1:100 |
| Goat anti-rabbit IgG (H+L), Alexa Fluor 488 | Goat | Invitrogen (Thermo Fisher Scientific) | A11034 | 1:1000 |
| Goat anti-mouse IgG (H+L), Alexa Fluor 568 | Goat | Invitrogen (Thermo Fisher Scientific) | A11004 | 1:1000 |
| Goat anti-rabbit IgG (H+L), Alexa Fluor 635 | Goat | Invitrogen (Thermo Fisher Scientific) | A31576 | 1:1000 |
| Goat anti-mouse IgG (H+L), Alexa Fluor 488 | Goat | Invitrogen (Thermo Fisher Scientific) | A11001 | 1:1000 |
| Goat anti-rabbit IgG (H+L), Alexa Fluor 647 | Goat | Invitrogen (Thermo Fisher Scientific) | A 21244 | 1:1000 |
| Goat anti-rabbit IgG (H+L), Alexa Fluor 568 | Goat | Invitrogen (Thermo Fisher Scientific) | A 11011 | 1:1000 |
| Goat anti-mouse IgG (H+L), Alexa Fluor 647 | Goat | Invitrogen (Thermo Fisher Scientific) | A 21235 | 1:1000 |
| Donkey anti-goat IgG (H+L), Alexa Fluor 647 | Donkey | Invitrogen (Thermo Fisher Scientific) | A21447 | 1:1000 |
| Donkey anti-goat IgG (H+L), Alexa Fluor 488 | Donkey | Invitrogen (Thermo Fisher Scientific) | A11055 | 1:1000 |
| Donkey anti-goat IgG (H+L), Alexa Fluor 568 | Donkey | Invitrogen (Thermo Fisher Scientific) | A11057 | 1:1000 |

### Whole mount staining and clearing

The staining and clearing protocols were optimized for inner ear organoids based on previously published methods (Park et al., 2019; van der Valk et al., 2026). IEOs were collected, rinsed once with PBS and fixed with 4% PFA for 20 min at room temperature. Samples were washed three times with PBS and then cleared using SHIELD (LifeCanvas Technologies). Samples were immersed in SHIELD-OFF, consisting of 25% deionised water, 25% SHIELD buffer and 50% SHIELD epoxy, with a maximum of three aggregates per microcentrifuge tube. On day 4, samples were transferred to the SHIELD-ON solution (include 12.5% Shield epoxy and 87.5% Shield-on buffer) for 24 hours at room temperature. After the SHIELD-ON treatment, samples were placed into a delipidation solution and incubated 24h at 37°C. Subsequently, immunohistochemistry was carried out. Aggregates were washed three times in PBST2 (0.2% TritonX-100 in PBS) for 10 minutes at room temperature. They were then blocked in 10% goat serum in PBST2 at room temperature for 1.5h, incubated in primary antibodies in 5% goat serum in PBST2 and rotated overnight at room temperature. Samples were washed with PBST2 five times for 10 minutes each and incubated with secondary antibodies in 5% goat serum in PBST2 on a rotator at room temperature overnight. Samples were washed with PBST2 five times for 10 minutes each before being incubated with DAPI (1 µg/mL in PBST2) for 4hours at room temperature. Aggregates were then rinsed with PBS 5 times for 10 minutes each and incubated in EasyIndex solution overnight at room temperature.

### Sample preparation for light sheet imaging

For light sheet imaging, aggregates were submerged in 1% low melting point agarose (Promega, cat. V2111) in EasyIndex, loaded into glass capillaries (Brand, cat. 701908, 701910) using the corresponding plungers (Brand, cat. 701934, 701936). Samples were stored at 4°C overnight, before being imaged using Zeiss light sheet Z.1 microscope.

### 3D image processing and analysis

3D image analysis was performed using Imaris (version 10.2.0) on light-sheet datasets acquired from individual IEOs. Multichannel image stacks were examined using both 3D view and serial slice visualization modes to enable three-dimensional inspection of OV structures.

Otic vesicles were identified based on predefined morphological and marker-based criteria. Morphologically, OVs were defined as spheroid or quasi-spherical epithelial vesicles containing a central lumen in 3D view. In serial slices view, as ring-like epithelial structure with a hollow centre and a regular, radially arranged nuclear pattern, as visualized by DAPI nuclear staining. In addition, OV identity required positive expression of at least one OV marker.

To count OV numbers within each aggregate, serial slices were examined throughout the entire z-stack. A structure was counted as OV, when a continuous vesicular structure was observed, beginning with the initial appearance of a localised epithelial cell cluster (“patch”), progressing through a well-defined ring-shaped vesicle, until the structure was no longer present.

To quantify OV size, the longest axis of each vesicle was measured. For each OV, linear measurements were obtained independently along the X, Y, and Z axes using Imaris measurement tools. Each axis was measured three times, and the maximum value among all measurements was recorded as the longest axis for that vesicle.

Marker expression patterns within OVs were assessed based on their spatial distribution in both 3D views and serial slices. For individual OVs with detectable marker expression, the signal was considered to display a spatial pattern when a non-uniform distribution was observed within the vesicle, including regionally localized expression (e.g., HMX3; Fig. 4A) or a graded spatial distribution across the OV (e.g., PAX2; Fig. 4A).

### 3D surface rendering and spatial orientation analysis of marker expression

For 3D surface rendering and segmentation of OVs were performed using the Surface module in Imaris. A machine learning–based approach was applied to segment OVs from the surrounding background. For training, OVs and background regions were manually annotated in at least 100 annotations to train the segmentation algorithm. The segmentation results were manually inspected in serial slices to confirm OV identity. Only surfaces corresponding to OVs that met the predefined identification criteria were retained for subsequent analysis. Surface rendered OVs were shown using colour coding based on object size for visualisation purposes. A continuous colour scale was applied to map surface volume values to colour, as shown in the corresponding scale bar.

For each segmented OV, the 3D spatial centre coordinates (X, Y, Z) were extracted using the Surface statistics function in Imaris. The OV centre was defined as the centre of homogeneous mass provided in the detailed statistics output and was recorded for downstream analyses. To delineate marker expression domains, background subtraction was applied to each marker channel using the Surface module. Marker-specific surfaces were generated using automatic intensity thresholding, followed by minor manual adjustments where necessary. The same threshold was applied to all OVs for each marker within an IEO. The resulting surfaces were inspected in both 3D and serial slice views to confirm their localisation within individual OVs. Only surfaces corresponding to OV associated expression patterns were retained. The centre of each marker expression domain was determined using the same centre of homogeneous mass parameter.

For each marker in each OV, a vector was calculated from the OV centre to the centre of the marker expression domain, for patterned OVs only. Centres were extracted as the centre of homogeneous mass of each surface i.e. the geometric centroid, not weighted by signal intensity. Marker domains were defined as above, expression outside the OV epithelium was excluded. Vectors were calculated in MATLAB (R2024) in the coordinate frame of the light-sheet acquisition and normalised to unit length (uᵢ), so that they carry direction only. Vectors were analysed separately for each marker within each IEO. Because OVs within an IEO are not independent, these analyses are descriptive and no statistical tests were applied.

To assess the overall alignment of marker expression domains, the mean resultant length was calculated as R = ǁ(1/n) Σ uᵢǁ, where n is the number of OVs. R ranges from 0 (no net alignment) to 1 (all vectors pointing in the same direction). As R depends on n, reference distributions were generated using Monte Carlo simulation. For each sample size, n independent unit vectors with uniformly random directions in three-dimensional space were generated, and R was calculated. This procedure was repeated 2,000,000 times. The mean and central 95% range (2.5th to 97.5th percentiles) of the simulated R values were used as descriptive references for comparison with the observed R.

Cosine similarity between individual OVs was calculated as the dot product of their unit vectors for every unique pair of OVs within an IEO. A cosine similarity of 1 indicates that two vectors point in the same direction, 0 indicates perpendicular directions and −1 indicates opposite directions. The angle (θ) between each pair of vectors was calculated as the inverse cosine (arccos) of the cosine similarity, and the results were displayed as heatmaps. For descriptive analysis, OV pairs were classified as similar (0° ≤ θ ≤ 45°), intermediate (45° < θ < 135°) or approximately opposite (135° ≤ θ ≤ 180°). These thresholds were chosen as the midpoints between identical (0°), perpendicular (90°) and opposite (180°) directions. For randomly oriented 3D vectors, 14.6%, 70.7% and 14.6% of pairs are expected in these categories. Each pair was counted once, and percentages were calculated using the total number of pairs for the marker in that IEO.

To assess the orientation of marker expression domains relative to the IEO centre, a 3D surface encompassing the entire IEO was generated from the DAPI channel in Imaris. The IEO centre was defined as the centre of homogeneous mass of this surface. For each OV, a reference vector was defined from the IEO centre to the OV centre. The angle between this reference vector and the corresponding marker expression vector was calculated in three dimensions. Marker vectors forming an angle of less than 90° with the reference vector were classified as outward, as they contained a component directed away from the IEO centre, whereas vectors forming an angle greater than 90° were classified as inward. Vectors perpendicular to the reference vector (90°) had neither an inward nor an outward component. Thus, outward orientation did not require exact alignment with the reference vector. For randomly oriented vectors, 50% are expected to be outward. For visualisation, marker expression vectors were translated to a common origin while preserving their original directions. Outward and inward vectors were represented by solid and dashed lines, respectively. The numbers of outward and inward vectors were recorded for each marker within each IEO.

### Statistical analysis

Statistical analyses were performed using GraphPad Prism (version 10.6.1) and MATLAB (R2024). Data were obtained from three independent differentiation experiments. Individual IEO aggregates were treated as independent biological replicates for all statistical analyses. OV measurements were summarized at the aggregate level, including OV number, median OV size and the proportions of OVs displaying marker expression or spatial patterning. The vector-based analyses were performed on a single IEO per timepoint and are descriptive; no statistical tests were applied.

Normality was assessed using the Shapiro–Wilk test. Depending on data distribution and experimental design, two-sided Mann–Whitney U tests, Welch’s t-tests, paired Wilcoxon signed-rank tests, or Kruskal–Wallis tests followed by Dunn’s multiple comparisons were used. P values were adjusted using the Benjamini–Hochberg false discovery rate or Bonferroni correction, as detailed in Supplementary Table 6. Adjusted P < 0.05 was considered statistically significant.

The 3D vector-based orientation analyses were descriptive, as only one IEO was analysed per time point. No statistical significance tests were performed. Monte Carlo simulations (2,000,000 iterations) of uniformly random 3D unit vectors were used to generate reference distributions for mean resultant length (R), with the mean and central 95% range used for descriptive comparison.

## Data availability

We have re-used publish scRNAseq datasets from (Steinhart et al., 2023); the analysis pipeline is available at https://github.com/Streit-lab/Organoid_OV_spatial_patterning.

## Author contributions

AGU optimised the IEO protocol, generated the PAX2 reporter line together with GC, and carried out its characterisation. AGU and JY performed all IEO cultures and 2D marker analysis. JY and MA developed IEO clearing protocols and light sheet imaging. JY performed all 3D data acquisition and analysis. AT and DJP carried out bioinformatics analysis. AGU and JY generated all figures; AS wrote the manuscript together with AGU. TZ generated OV patterning schematics and MATLAB vector visualisations. All authors commented and edited the manuscript. AS obtained funding, supervised and directed the project.

## Acknowledgements

The authors are grateful to Mandar Phatak for expert advice on imaging and image analysis, Anbo Dong for advice and assistance with statistical analysis, Yuhan Lu for advice and assistance with OV orientation analysis, and Robert C. Hayward, Susmitha Rao and Giang Duong for excellent technical support. We thank Karl Koehler for advice and sharing their IEO protocol prior to publication, Claudio D. Stern and Zoe Mann for comments on the manuscript. This work was supported by the BBSRC-funded London Interdisciplinary Biosciences Programme (LIDo) through an iCase studentship co-funded by Boehringer Ingelheim and King’s Doctoral College, and by BBSRC project grants BB/V006290/1 and BB/V006339/1 to AS.

## References

Alsina, B., Giraldez, F., Pujades, C., 2009. Patterning and cell fate in ear development. Int J Dev Biol 53, 1503–1513.

Barald, K.F., Kelley, M.W., 2004. From placode to polarization: new tunes in inner ear development. Development 131, 4119–4130.

Bhaduri, A., Andrews, M.G., Kriegstein, A.R., Nowakowski, T.J., 2020a. Are Organoids Ready for Prime Time? Cell Stem Cell 27, 361–365.

Bhaduri, A., Andrews, M.G., Mancia Leon, W., Jung, D., Shin, D., Allen, D., Jung, D., Schmunk, G., Haeussler, M., Salma, J., Pollen, A.A., Nowakowski, T.J., Kriegstein, A.R., 2020b. Cell stress in cortical organoids impairs molecular subtype specification. Nature 578, 142–148.

Bok, J., Bronner-Fraser, M., Wu, D.K., 2005. Role of the hindbrain in dorsoventral but not anteroposterior axial specification of the inner ear. Development 132, 2115–2124.

Bok, J., Chang, W., Wu, D.K., 2007a. Patterning and morphogenesis of the vertebrate inner ear. Int J Dev Biol 51, 521–533.

Bok, J., Dolson, D.K., Hill, P., Ruther, U., Epstein, D.J., Wu, D.K., 2007b. Opposing gradients of Gli repressor and activators mediate Shh signaling along the dorsoventral axis of the inner ear. Development 134, 1713–1722.

Bok, J., Raft, S., Kong, K.A., Koo, S.K., Drager, U.C., Wu, D.K., 2011. Transient retinoic acid signaling confers anterior-posterior polarity to the inner ear. Proc Natl Acad Sci U S A 108, 161–166.

Brigande, J.V., Kiernan, A.E., Gao, X., Iten, L.E., Fekete, D.M., 2000. Molecular genetics of pattern formation in the inner ear: Do compartment boundaries play a role? [In Process Citation]. Proc Natl Acad Sci U S A 97, 11700–11706.

Brugmann, S.A., Pandur, P.D., Kenyon, K.L., Pignoni, F., Moody, S.A., 2004. Six1 promotes a placodal fate within the lateral neurogenic ectoderm by functioning as both a transcriptional activator and repressor. Development 131, 5871–5881.

Chang, W., Lin, Z., Kulessa, H., Hebert, J., Hogan, B.L., Wu, D.K., 2008. Bmp4 is essential for the formation of the vestibular apparatus that detects angular head movements. PLoS Genet 4, e1000050.

Chiaradia, I., Imaz-Rosshandler, I., Nilges, B.S., Boulanger, J., Pellegrini, L., Das, R., Kashikar, N.D., Lancaster, M.A., 2023. Tissue morphology influences the temporal program of human brain organoid development. Cell Stem Cell 30, 1351–1367 e1310.

Chiaradia, I., Lancaster, M.A., 2026. Diversity and complexity in neural organoids. FEBS Lett.

Doda, D., Alonso Jimenez, S., Rehrauer, H., Carreno, J.F., Valsamides, V., Di Santo, S., Widmer, H.R., Edge, A., Locher, H., van der Valk, W.H., Zhang, J., Koehler, K.R., Roccio, M., 2023. Human pluripotent stem cell-derived inner ear organoids recapitulate otic development in vitro. Development 150.

Driver, E.C., Kelley, M.W., 2020. Development of the cochlea. Development 147.

Duncan, R.K., Liu, L., Moyer, M., Wylie, A., Dano, R., Cassinotti, L., 2025. Retinoic acid signaling guides the efficiency of inner ear organoid-genesis and governs sensory-nonsensory fate specification. Stem Cell Reports 20.

Groves, A.K., Bronner-Fraser, M., 2000. Competence, specification and commitment in otic placode induction. Development 127, 3489–3499.

Groves, A.K., Fekete, D.M., 2012. Shaping sound in space: the regulation of inner ear patterning. Development 139, 245–257.

Guo, J.Y., He, L., Chen, Z.R., Liu, K., Gong, S.S., Wang, G.P., 2021. AAV8-mediated Atoh1 overexpression induces dose-dependent regeneration of vestibular hair cells in adult mice. Neurosci Lett 747, 135679.

Hao, Y., Stuart, T., Kowalski, M.H., Choudhary, S., Hoffman, P., Hartman, A., Srivastava, A., Molla, G., Madad, S., Fernandez-Granda, C., Satija, R., 2024. Dictionary learning for integrative, multimodal and scalable single-cell analysis. Nature Biotechnology 42, 293–304.

Hidalgo-Sanchez, M., Alvarado-Mallart, R., Alvarez, I.S., 2000. Pax2, otx2, gbx2 and fgf8 expression in early otic vesicle development [In Process Citation]. Mech Dev 95, 225–229.

Hutson, M.R., Lewis, J.E., Nguyen-Luu, D., Lindberg, K.H., Barald, K.F., 1999. Expression of Pax2 and patterning of the chick inner ear. J Neurocytol 28, 795–807.

Jayasena, C.S., Ohyama, T., Segil, N., Groves, A.K., 2008. Notch signaling augments the canonical Wnt pathway to specify the size of the otic placode. Development 135, 2251–2261.

Kanton, S., Boyle, M.J., He, Z., Santel, M., Weigert, A., Sanchis-Calleja, F., Guijarro, P., Sidow, L., Fleck, J.S., Han, D., Qian, Z., Heide, M., Huttner, W.B., Khaitovich, P., Paabo, S., Treutlein, B., Camp, J.G., 2019. Organoid single-cell genomic atlas uncovers human-specific features of brain development. Nature 574, 418–422.

Kelley, M.W., 2022. Cochlear Development; New Tools and Approaches. Front Cell Dev Biol 10, 884240.

Kelly, M.C., Chang, Q., Pan, A., Lin, X., Chen, P., 2012. Atoh1 directs the formation of sensory mosaics and induces cell proliferation in the postnatal mammalian cochlea in vivo. J Neurosci 32, 6699–6710.

Koehler, K.R., Nie, J., Longworth-Mills, E., Liu, X.P., Lee, J., Holt, J.R., Hashino, E., 2017. Generation of inner ear organoids containing functional hair cells from human pluripotent stem cells. Nat Biotechnol 35, 583–589.

Lancaster, M.A., Knoblich, J.A., 2014a. Generation of cerebral organoids from human pluripotent stem cells. Nat Protoc 9, 2329–2340.

Lancaster, M.A., Knoblich, J.A., 2014b. Organogenesis in a dish: modeling development and disease using organoid technologies. Science 345, 1247125.

Lawoko-Kerali, G., Rivolta, M.N., Holley, M., 2002. Expression of the transcription factors GATA3 and Pax2 during development of the mammalian inner ear. J Comp Neurol 442, 378–391.

Lin, Z., Cantos, R., Patente, M., Wu, D.K., 2005. Gbx2 is required for the morphogenesis of the mouse inner ear: a downstream candidate of hindbrain signaling. Development 132, 2309–2318.

Lin, Z., Perez, P., Sun, Z., Liu, J.J., Shin, J.H., Hyrc, K.L., Samways, D., Egan, T., Holley, M.C., Bao, J., 2012. Reprogramming of single-cell-derived mesenchymal stem cells into hair cell-like cells. Otol Neurotol 33, 1648–1655.

Litsiou, A., Hanson, S., Streit, A., 2005. A balance of FGF, Wnt and BMP signalling positions the future placode territory in the head. Development 132, 4051–4062.

Liu, X.P., Koehler, K.R., Mikosz, A.M., Hashino, E., Holt, J.R., 2016. Functional development of mechanosensitive hair cells in stem cell-derived organoids parallels native vestibular hair cells. Nat Commun 7, 11508.

Liu, Z., Dearman, J.A., Cox, B.C., Walters, B.J., Zhang, L., Ayrault, O., Zindy, F., Gan, L., Roussel, M.F., Zuo, J., 2012. Age-dependent in vivo conversion of mouse cochlear pillar and Deiters’ cells to immature hair cells by Atoh1 ectopic expression. J Neurosci 32, 6600–6610.

Luo, T., Liu, C., Cheng, T., Zhao, G.-Q., Huang, Y., Luan, J.-Y., Guo, J., Liu, X., Wang, Y.-F., Dong, Y., Xiao, Y., He, E., Sun, R.-Z., Chen, X., Chen, J., Ma, J., Megason, S., Ji, J., Xu, P.-F., 2025. Establishing dorsal-ventral patterning in human neural tube organoids with synthetic organizers. Cell Stem Cell 32, 1071–1086.e1078.

Marsh, S.E., 2021. scCustomize: Custom Visualizations & Functions for Streamlined Analyses of Single Cell Sequencing.

Merlo, G.R., Paleari, L., Mantero, S., Zerega, B., Adamska, M., Rinkwitz, S., Bober, E., Levi, G., 2002. The Dlx5 homeobox gene is essential for vestibular morphogenesis in the mouse embryo through a BMP4-mediated pathway. Dev Biol 248, 157–169.

Moore, S.T., Nakamura, T., Nie, J., Solivais, A.J., Aristizabal-Ramirez, I., Ueda, Y., Manikandan, M., Reddy, V.S., Romano, D.R., Hoffman, J.R., Perrin, B.J., Nelson, R.F., Frolenkov, G.I., Chuva de Sousa Lopes, S.M., Hashino, E., 2023. Generating high-fidelity cochlear organoids from human pluripotent stem cells. Cell Stem Cell 30, 950–961 e957.

Murtazina, A., Fatieieva, Y., Waern, F., Maunsell, H.R., Thawani, A., Semsch, B., Bostrom, J., Reagor, C.C., Kameneva, P., Araslanova, K., Isaev, S., Schelb, F., Fried, K., Erickson, A.G., Klimovich, A., Streit, A., Kutscher, L.M., Kozmikova, I., Kozmik, Z., Andersson, E.R., Schlosser, G., Groves, A.K., Adameyko, I., 2026. Single-cell, clonal and spatial atlases of cranial placodes illuminate their specification and evolution. bioRxiv.

Nakajima, Y., 2015. Signaling regulating inner ear development: cell fate determination, patterning, morphogenesis, and defects. Congenit Anom (Kyoto) 55, 17–25.

Nist-Lund, C., Kim, J., Koehler, K.R., 2022. Advancements in inner ear development, regeneration, and repair through otic organoids. Curr Opin Genet Dev 76, 101954.

Ohlenschlaeger, M.S., Jensen, P., Havelund, J.F., Schmidt, S.I., Mohamed, F.A., Sutcliffe, M., Elmkvist, S.B., Criscuolo, L., Wingett, S.W., Chiaradia, I., Bayram, E., Nicolaisen, J.A.A., Jakobsen, L.A., Brewer, J., Benros, M.E., Freude, K., Faergeman, N.J., Lancaster, M.A., Larsen, M.R., Bogetofte, H., 2026. Multi-omic analysis of guided and unguided forebrain organoids reveals differences in cellular composition and metabolic profiles. Cell Rep Methods 6, 101295.

Ohta, S., Mansour, S.L., Schoenwolf, G.C., 2010. BMP/SMAD signaling regulates the cell behaviors that drive the initial dorsal-specific regional morphogenesis of the otocyst. Dev Biol 347, 369–381.

Ohta, S., Schoenwolf, G.C., 2018. Hearing crosstalk: the molecular conversation orchestrating inner ear dorsoventral patterning. Wiley Interdiscip Rev Dev Biol 7.

Ohta, S., Wang, B., Mansour, S.L., Schoenwolf, G.C., 2016a. BMP regulates regional gene expression in the dorsal otocyst through canonical and non-canonical intracellular pathways. Development 143, 2228–2237.

Ohta, S., Wang, B., Mansour, S.L., Schoenwolf, G.C., 2016b. SHH ventralizes the otocyst by maintaining basal PKA activity and regulating GLI3 signaling. Dev Biol 420, 100–109.

Ohyama, T., Mohamed, O.A., Taketo, M.M., Dufort, D., Groves, A.K., 2006. Wnt signals mediate a fate decision between otic placode and epidermis. Development 133, 865–875.

Park, Y.-G., Sohn, C.H., Chen, R., McCue, M., Yun, D.H., Drummond, G.T., Ku, T., Evans, N.B., Oak, H.C., Trieu, W., Choi, H., Jin, X., Lilascharoen, V., Wang, J., Truttmann, M.C., Qi, H.W., Ploegh, H.L., Golub, T.R., Chen, S.-C., Frosch, M.P., Kulik, H.J., Lim, B.K., Chung, K., 2019. Protection of tissue physicochemical properties using polyfunctional crosslinkers. Nature Biotechnology 37, 73–83.

Pianigiani, G., Roccio, M., 2024. Inner Ear Organoids: Strengths and Limitations. J Assoc Res Otolaryngol 25, 5–11.

Riccomagno, M.M., Martinu, L., Mulheisen, M., Wu, D.K., Epstein, D.J., 2002. Specification of the mammalian cochlea is dependent on Sonic hedgehog. Genes Dev 16, 2365–2378.

Riccomagno, M.M., Takada, S., Epstein, D.J., 2005. Wnt-dependent regulation of inner ear morphogenesis is balanced by the opposing and supporting roles of Shh. Genes Dev 19, 1612–1623.

Rifes, P., Isaksson, M., Rathore, G.S., Aldrin-Kirk, P., Møller, O.K., Barzaghi, G., Lee, J., Egerod, K.L., Rausch, D.M., Parmar, M., Pers, T.H., Laurell, T., Kirkeby, A., 2020. Modeling neural tube development by differentiation of human embryonic stem cells in a microfluidic WNT gradient. Nat Biotechnol 38, 1265–1273.

Rossi, G., Manfrin, A., Lutolf, M.P., 2018. Progress and potential in organoid research. Nat Rev Genet 19, 671–687.

Rumbo, M., Alsina, B., 2024. Cellular diversity of human inner ear organoids revealed by single-cell transcriptomics. Development 151.

Sampath Kumar, A., Tian, L., Bolondi, A., Hernández, A.A., Stickels, R., Kretzmer, H., Murray, E., Wittler, L., Walther, M., Barakat, G., Haut, L., Elkabetz, Y., Macosko, E.Z., Guignard, L., Chen, F., Meissner, A., 2023. Spatiotemporal transcriptomic maps of whole mouse embryos at the onset of organogenesis. Nat Genet 55, 1176–1185.

Soldatov, R., Kaucka, M., Kastriti, M.E., Petersen, J., Chontorotzea, T., Englmaier, L., Akkuratova, N., Yang, Y., Häring, M., Dyachuk, V., Bock, C., Farlik, M., Piacentino, M.L., Boismoreau, F., Hilscher, M.M., Yokota, C., Qian, X., Nilsson, M., Bronner, M.E., Croci, L., Hsiao, W.-Y., Guertin, D.A., Brunet, J.-F., Consalez, G.G., Ernfors, P., Fried, K., Kharchenko, P.V., Adameyko, I., 2019. Spatiotemporal structure of cell fate decisions in murine neural crest. Science 364, eaas9536.

Staecker, H., Praetorius, M., Baker, K., Brough, D.E., 2007. Vestibular hair cell regeneration and restoration of balance function induced by math1 gene transfer. Otol Neurotol 28, 223–231.

Steinhart, M.R., van der Valk, W.H., Osorio, D., Serdy, S.A., Zhang, J., Nist-Lund, C., Kim, J., Moncada-Reid, C., Sun, L., Lee, J., Koehler, K.R., 2023. Mapping oto-pharyngeal development in a human inner ear organoid model. Development 150.

Streit, A., 2002. Extensive cell movements accompany formation of the otic placode. Dev Biol 249, 237–254.

Subkhankulova, T., Camargo Sosa, K., Uroshlev, L.A., Nikaido, M., Shriever, N., Kasianov, A.S., Yang, X., Rodrigues, F., Carney, T.J., Bavister, G., Schwetlick, H., Dawes, J.H.P., Rocco, A., Makeev, V.J., Kelsh, R.N., 2023. Zebrafish pigment cells develop directly from persistent highly multipotent progenitors. Nat Commun 14, 1258.

Tang, P.C., Alex, A.L., Nie, J., Lee, J., Roth, A.A., Booth, K.T., Koehler, K.R., Hashino, E., Nelson, R.F., 2019. Defective Tmprss3-Associated Hair Cell Degeneration in Inner Ear Organoids. Stem Cell Reports 13, 147–162.

Thiery, A.P., Buzzi, A.L., Hamrud, E., Cheshire, C., Luscombe, N.M., Briscoe, J., Streit, A., 2023. scRNA-sequencing in chick suggests a probabilistic model for cell fate allocation at the neural plate border. Elife 12.

Torres, M., Gomez-Pardo, E., Gruss, P., 1996. Pax2 contributes to inner ear patterning and optic nerve trajectory. Development 122, 3381–3391.

Ueda, Y., Nakamura, T., Nie, J., Solivais, A.J., Hoffman, J.R., Daye, B.J., Hashino, E., 2023. Defining developmental trajectories of prosensory cells in human inner ear organoids at single-cell resolution. Development 150.

van der Valk, W.H., Nist-Lund, C., Zhang, J., Perea, C., Jin, J., Gim, K.Y., Steinhart, M.R., Lee, J., Koehler, K.R., 2026. Generation and characterization of vestibular inner ear organoids from human pluripotent stem cells. Nat Protoc 21, 391–428.

van der Valk, W.H., Steinhart, M.R., Zhang, J., Koehler, K.R., 2021. Building inner ears: recent advances and future challenges for in vitro organoid systems. Cell death and differentiation 28, 24–34.

van der Valk, W.H., van Beelen, E.S.A., Steinhart, M.R., Nist-Lund, C., Osorio, D., de Groot, J., Sun, L., van Benthem, P.P.G., Koehler, K.R., Locher, H., 2023. A single-cell level comparison of human inner ear organoids with the human cochlea and vestibular organs. Cell Rep 42, 113527.

Velasco, S., Kedaigle, A.J., Simmons, S.K., Nash, A., Rocha, M., Quadrato, G., Paulsen, B., Nguyen, L., Adiconis, X., Regev, A., Levin, J.Z., Arlotta, P., 2019. Individual brain organoids reproducibly form cell diversity of the human cerebral cortex. Nature 570, 523–527.

Wang, T., Yang, T., Kedaigle, A., Pregernig, G., McCarthy, R., Holmes, B., Wu, X., Becker, L., Pan, N., So, K., Chen, L., He, J., Mahmoudi, A., Negi, S., Kowalczyk, M., Gibson, T., Druckenbrod, N., Cheng, A.G., Burns, J., 2024. Precise genetic control of ATOH1 enhances maturation of regenerated hair cells in the mature mouse utricle. Nat Commun 15, 9166.

Warchol, M.E., Richardson, G.P., 2009. Expression of the Pax2 transcription factor is associated with vestibular phenotype in the avian inner ear. Dev Neurobiol 69, 191–202.

Whitfield, T.T., Hammond, K.L., 2007. Axial patterning in the developing vertebrate inner ear. Int J Dev Biol 51, 507–520.

WorldHealthOrganisation, 2021. World Report on Hearing.

Wu, D.K., Kelley, M.W., 2012. Molecular mechanisms of inner ear development. Cold Spring Harbor perspectives in biology 4, a008409.

Xue, X., Kim, Y.S., Ponce-Arias, A.I., O’Laughlin, R., Yan, R.Z., Kobayashi, N., Tshuva, R.Y., Tsai, Y.H., Sun, S., Zheng, Y., Liu, Y., Wong, F.C.K., Surani, A., Spence, J.R., Song, H., Ming, G.L., Reiner, O., Fu, J., 2024. A patterned human neural tube model using microfluidic gradients. Nature 628, 391–399.

Xue, X., Rahman, O.M., Sun, S., Bok, J., Tang, A., Fu, J., 2026. Generation of spatially patterned human neural tube-like structures using microfluidic gradient devices. Nat Protoc 21, 1730–1753.

Zappia, L., Oshlack, A., 2018. Clustering trees: a visualization for evaluating clusterings at multiple resolutions. Gigascience 7.

Zou, D., Silvius, D., Rodrigo-Blomqvist, S., Enerback, S., Xu, P.-X., 2006. Eya1 regulates the growth of otic epithelium and interacts with Pax2 during the development of all sensory areas in the inner ear. Developmental Biology 298, 430–441.

