## Supplementary Figures for "Incomplete Spatial Patterning of Otic Vesicles in Human Inner Ear Organoids"

### Supplementary material

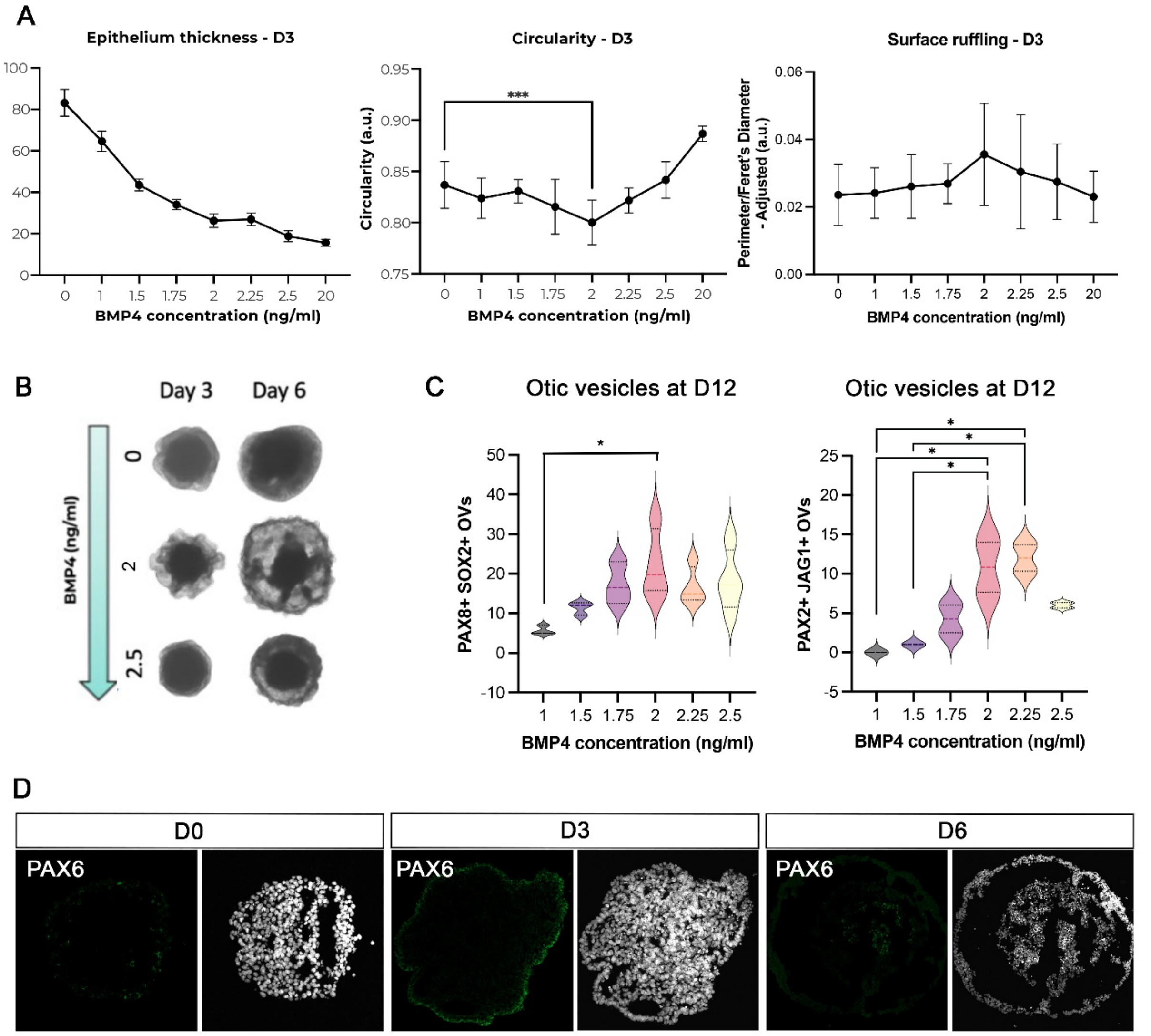


**Supplementary Figure 1. Titration of BMP4**
**A.** Morphological features of aggregates measured at day 3 (mean±s.d.). Epithelium thickness was measured at three different locations per aggregate. All BMP4 treatment concentrations differed significantly from control concentrations 0 and 20ng/ml (not shown). For circularity, only aggregates treated with 2ng/ml of BMP4 showed a statistically significant difference compared to 0ng/ml. All BMP4 treatment concentrations differed significantly from 20 ng/ml (not shown). None of the surface ruffling measurements showed statistically significant difference with any of the controls. All parameters were measured in 10 aggregates across three independent differentiations (n=3, 30 aggregates total). **B.** Representative images showing aggregate morphology following treatment with increasing BMP4 concentration. C. Number of vesicles positive for otic markers PAX8 and SOX2, and PAX2 and JAG1, respectively, at D12, assessed in 2-4 aggregates per differentiation across 3 independent differentiations (n =3) except for 1.75 ng/ml BMP4 concentration, which was assessed across only two independent differentiations (n=2). D. Representative images of organoids at D0, D3 and D6, stained for the neural marker PAX6 (left) and DAPI (right). Statistical analysis: Two-way ANOVA for A and mixed-effect model for C; significance levels are indicated as follows: $p< 0.05$ (*), $p<0.001$ (***) and $p<0.0001$ (****).

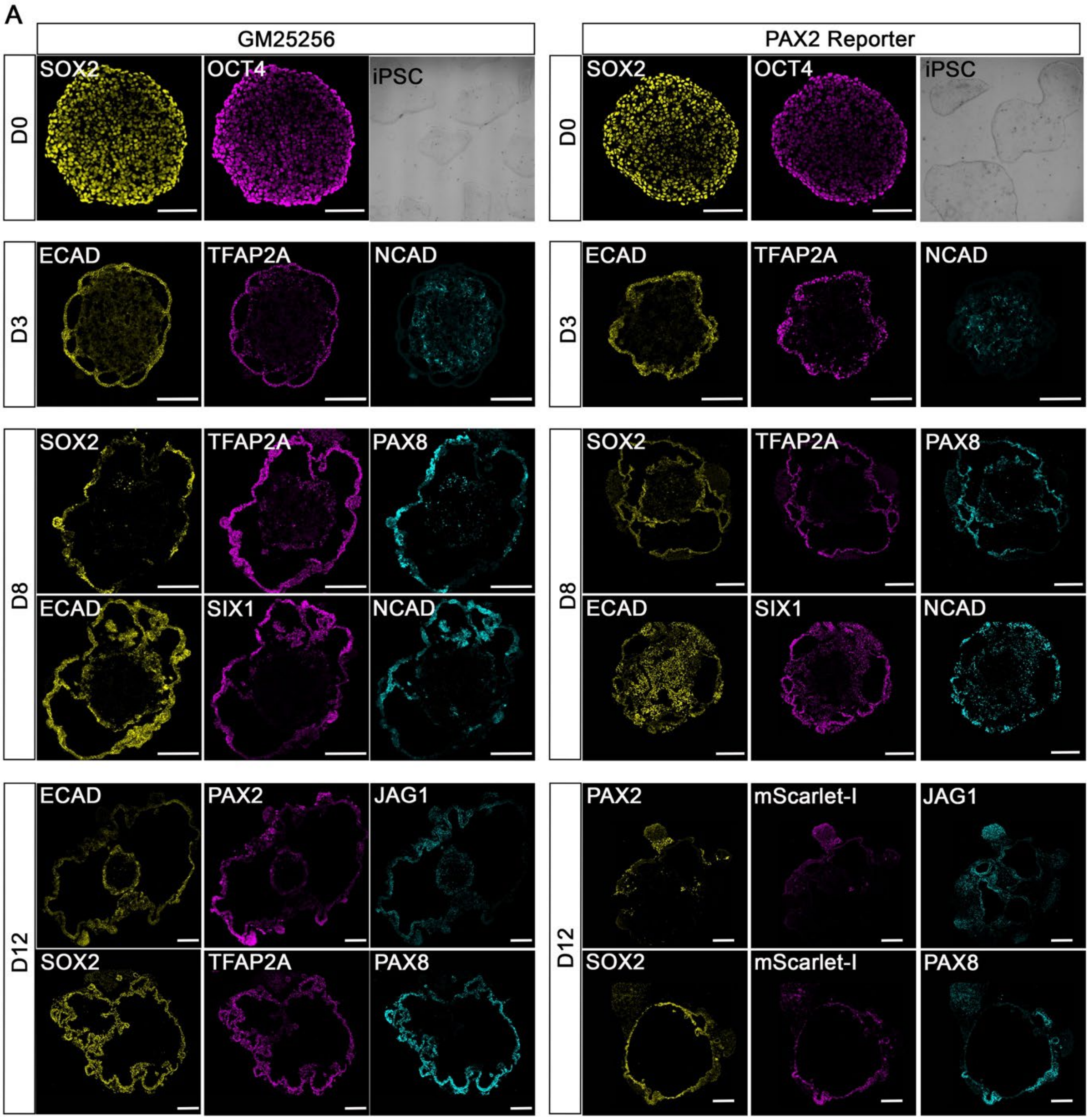


**Supplementary Figure 1. Characterisation of PAX2-P2A-mScarlet-I reporter cell line**

Expression of key otic markers during organoid differentiation and comparison with the parental line GM25256 confirmed otic induction in the new PAX2-P2A-mScarlet-I. For each cell line, marker expression at each time point was confirmed in 5–6 aggregates per differentiation. At least three independent differentiations per cell line were examined.

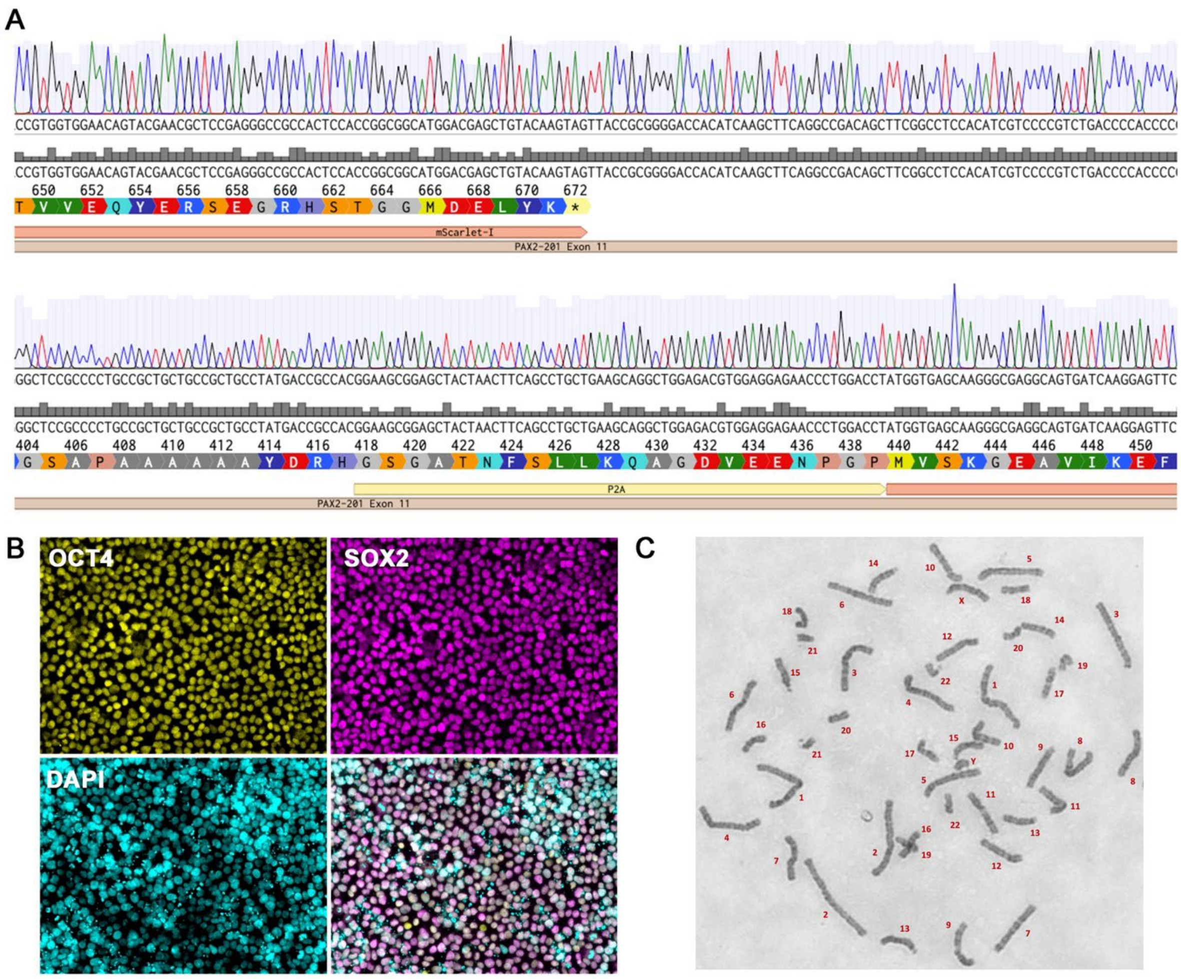


**Supplementary Figure 2**. **Quality check of PAX2-P2A-mScarlet-I reporter cell line confirms clean insertion of mScarlet-I.**

**A.** Sanger sequencing data showed correct monoallelic insertion of mScarlet-I, and a wild type allele free from indels. **B.** iPSC cultures show co-expression of pluripotency markers OCT4 and SOX2. **C**. Karyotype of PAX2-P2A-mScarlet-I reporter cell line. Chromosomes are organized and numbered in pairs (from 1 to 22), which represent the 22 autosomal chromosome pairs in humans. Chromosomes labelled ‘X’ and ‘Y’ are the sex chromosomes. No visible large deletions, duplications, or translocations, or extra or missing chromosomes are apparent.

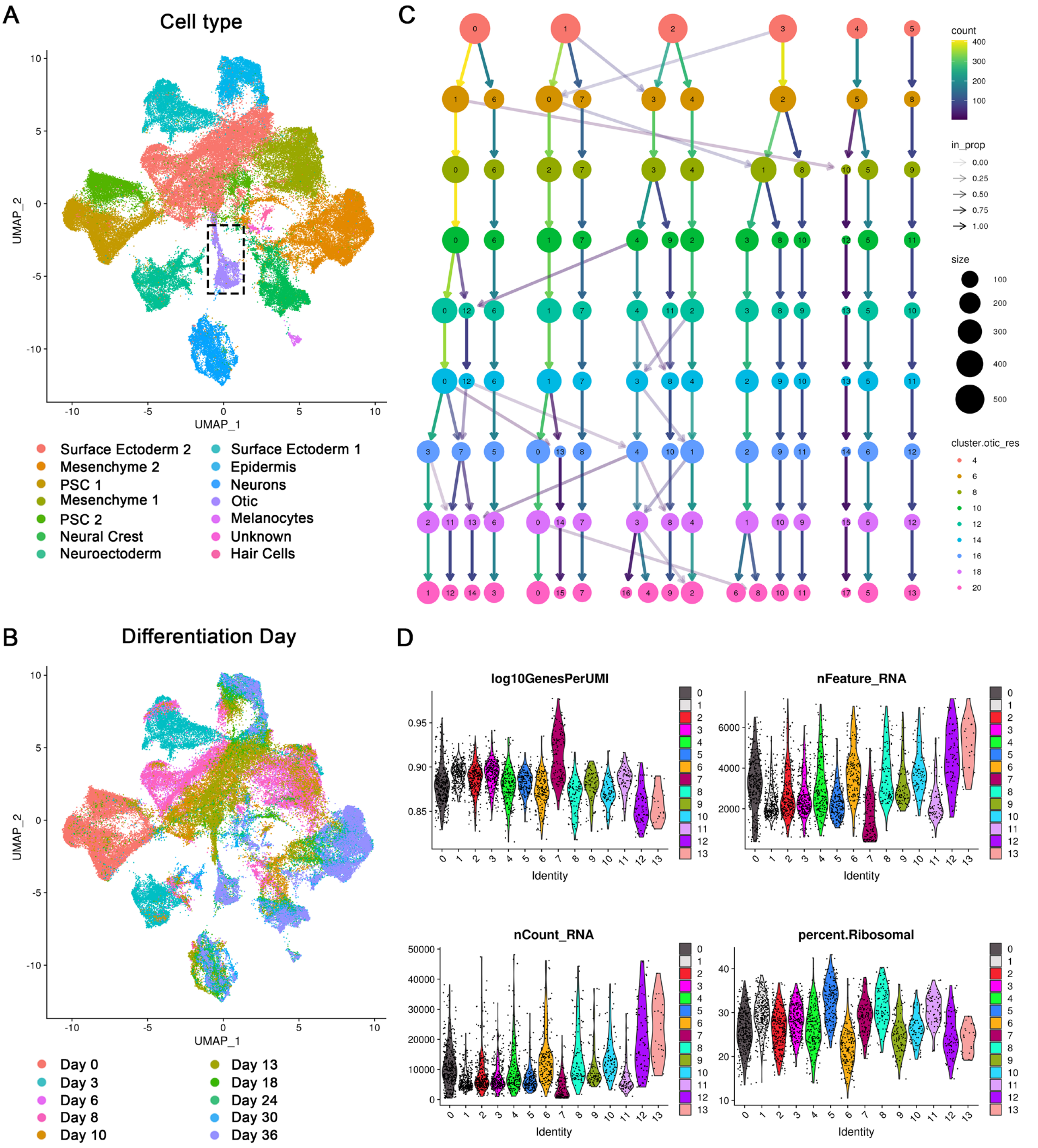


**Supplementary Figure 3. scRNAseq analysis from inner ear organoids.**
**A**. UMAP of scRNAseq data from IEOs (Steinhart et al., 2023): cell type identities were assigned to different cell clusters as downloaded from GSE188936. Boxed area represents the otic epithelium subset as defined in Steinhart et al, 2023 (purple). **B**. UMAP of aggregates from days 0-36 colour coded according to differentiation day. Note that the otic epithelium subset only contains cells from Day 18 onwards. **C.** Clustree analysis of clustering resolutions for the reclustered otic subset from Day 18-36 (Figure 2A). The number and identity of the clusters is relatively stable from a resolution of 1.2. **D.** QC characteristics of the clusters at resolution 1.2. While there is some variation in these parameters, particularly for smaller clusters such as 12 and 13, these were not out of the range of the other clusters. Note that the processed data from Steinhart, et al., 2023 did not contain any mitochondrial genes.

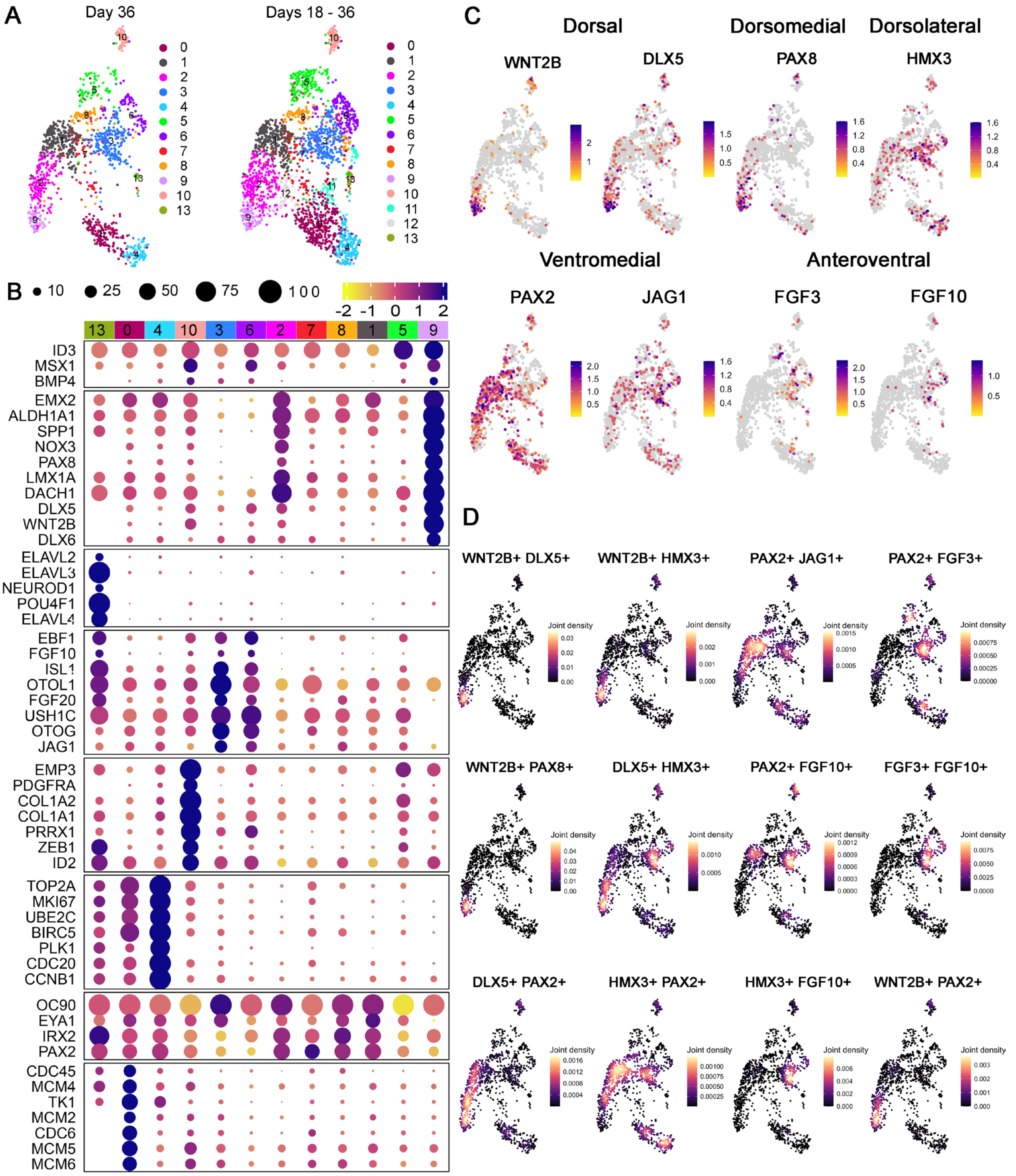


**Supplementary Figure 4. scRNAseq analysis of day 36 cells.**
**A.** The D18-36 otic subset was further subset using only the oldest cells (day 36). Comparison of the day 36 only cells (left) with the day 18-36 cells (right). Note that the day 36 cells show the similar distribution in the UMAP, except for the loss of clusters 11 and 12 which contained only earlier cells. Note: right image is identical to image in Figure 2A. **B.** Dotplot displaying the top differentially expressed genes in each cluster. Note the similarity of distribution of markers compared to Figure 2B. **C**. Feature plots showing the expression of known regional markers in OVs in vivo. These show the same distribution as Figure 2C. **D.** Joint density plots showing pairwise comparisons of the regional markers shown in C. Compare with the same plots shown in Figure 2D.

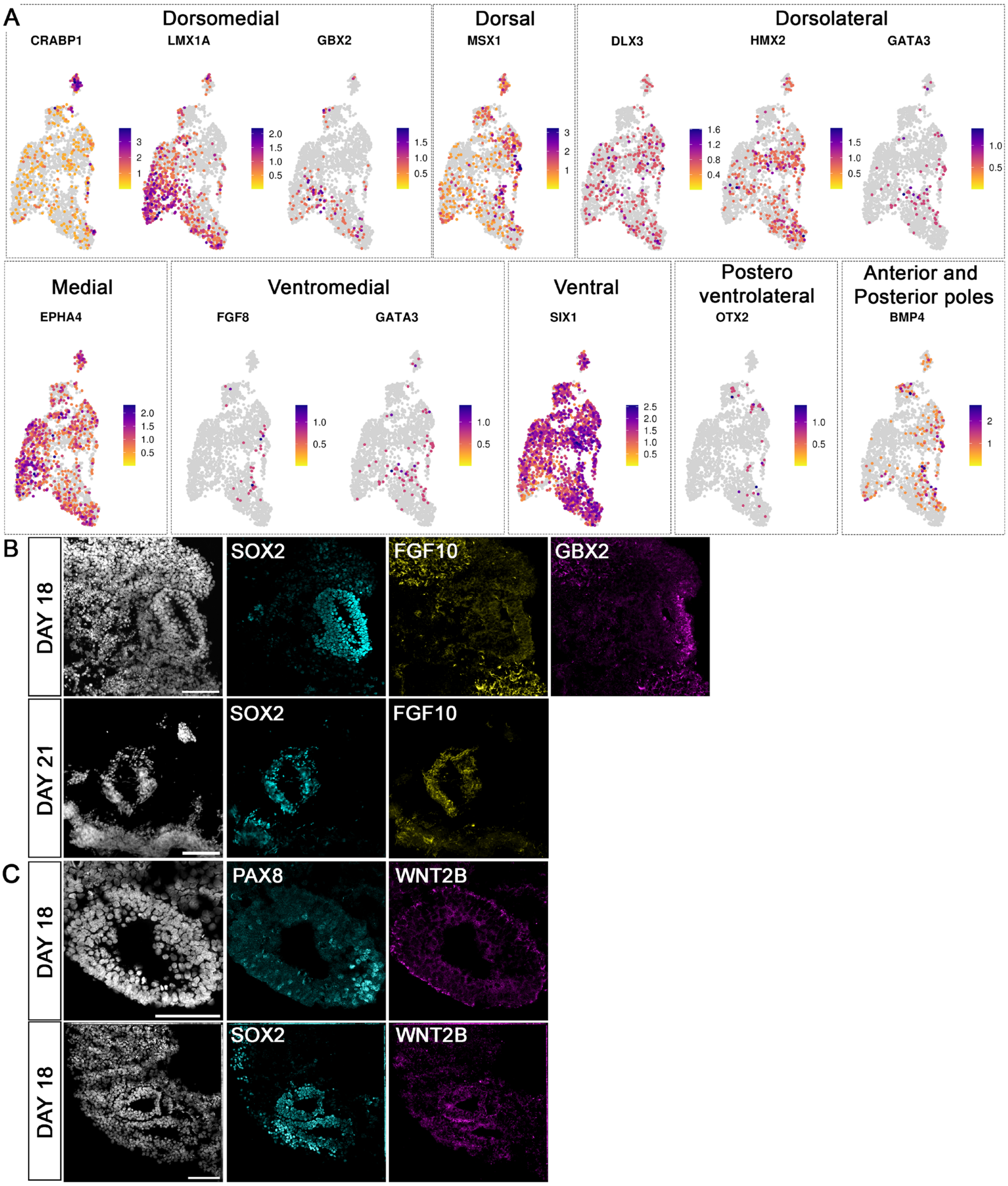


**Supplementary Figure 5. A.** Feature plots showing expression of key patterning genes. Note: the labels indicate the most representative location at otic vesicle stage. When chicken and mouse expression differ, mouse expression is shown in the label. B. At D18, FGF10 expressing cells surround the OV, which is positive for anteroventral SOX2 and dorsomedial GBX2 (patterned; n=6 in 2 independent differentiations). Co-expression of anteroventral markers FGF10 and SOX2 at D21 (n=5 2 independent differentiations). C. Expression of WNT2B at day 18 in the OV and surrounding cells. WNT2B is expressed in all cells (top row) or localised but co-expressed with SOX2 (n=7, 2 independent differentiations).

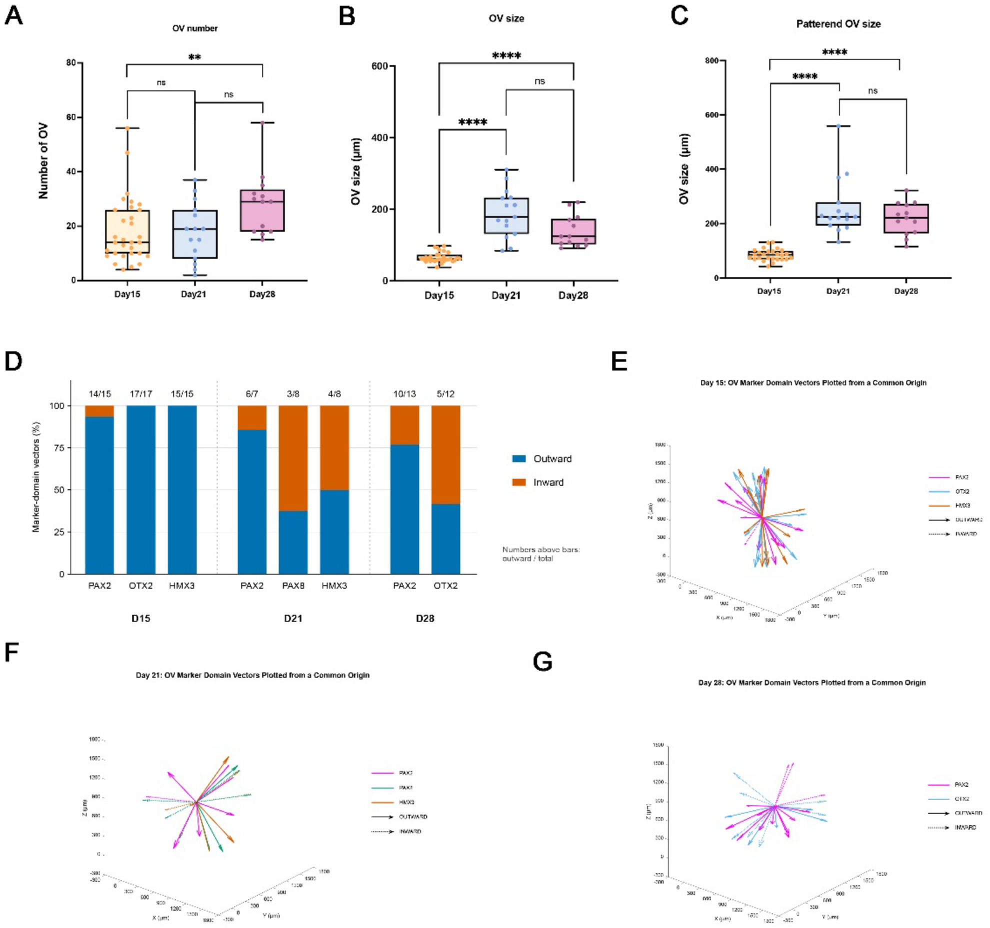


**Supplementary Figure 6. Quantitative analysis of otic vesicle number, size and spatial orientation of marker-expression domains in IEOs.**

**A.** Number of OVs per aggregate at D15 (n=31 IEOs), D21 (n=15) and D28 (n=13). Two-sided Mann–Whitney U tests (D15 vs D21, adjusted P=1.000; D15 vs D28, adjusted P=0.00812) or Welch's t-test (D21 vs D28, adjusted P=0.05923), with Bonferroni correction for three comparisons. **P < 0.01; ns, not significant. **B.** OV size, defined as the median longest OV diameter, at D15 (n=31), D21 (n=15) and D28 (n=13). Two-sided Mann–Whitney U tests (D15 vs D21, adjusted P=4.04 × $10^{-7}$; D15 vs D28, adjusted P=1.05 × $10^{-6}$) and Welch's t-test (D21 vs D28, adjusted P=0.104), with Bonferroni correction. **C**. Median diameter of patterned OVs per IEO at D15 (n=26), D21 (n=15) and D28 (n=13). Two-sided Mann–Whitney U tests (D15 vs D21, adjusted P=4.22 × $10^{-7}$; D21 vs D28, adjusted P=1.000) and Welch's t-test (D15 vs D28, adjusted P=7.16 × $10^{-6}$), with Bonferroni correction. Ns: not significant; * p-value < 0.05; ** p-value < 0.01; *** p-value < 0.001; **** p-value < 0.0001. **D.** Proportions of outward- and inward-oriented marker-domain vectors for each marker at D15, D21 and D28. Numbers above the bars indicate the number of outward vectors relative to the total analysed. **E–G.** 3D visualisation of marker domain vectors at D15 (E), D21 (F) and D28 (G). Outward and inward orientations were determined using the original positions of the OVs within the IEO before the marker vectors were moved to a common origin for visualisation in MATLAB. Colours indicate different markers, while solid and dashed arrows represent outward and inward vectors, respectively.

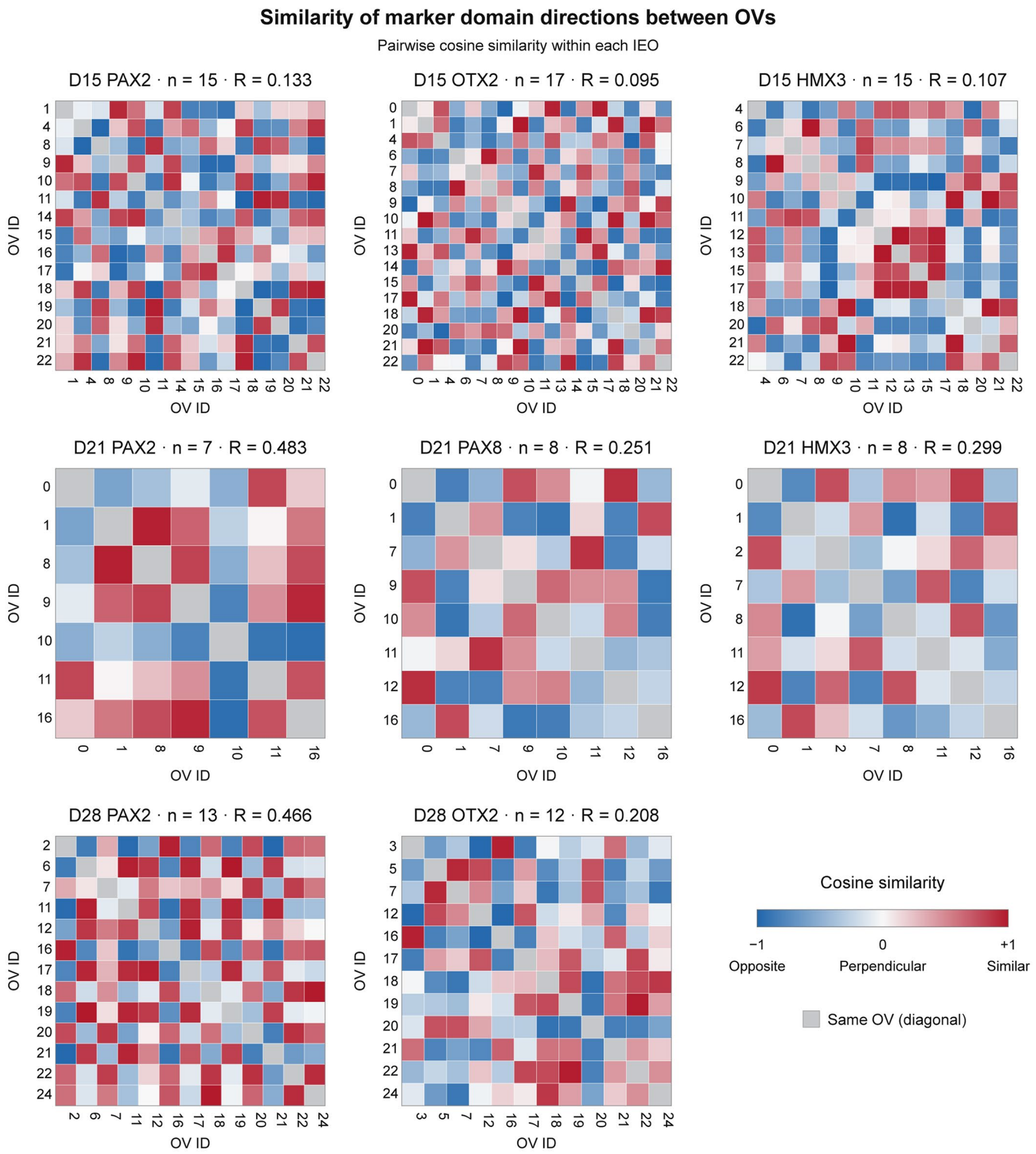


**Supplementary Figure 7. Orientation of marker expression domains across OVs within IEOs.**

The heatmaps show the cosine similarity of the directionality of each marker pair for pairwise comparison of all OVs in a single IEO. Red indicates similar directions (cosine similarity close to +1), white indicates approximately perpendicular directions (close to 0), and blue indicates opposite directions (close to −1). n denotes the number of OVs, and R denotes the mean resultant length, with higher values indicating greater overall alignment. One IEO was analysed at each time point.

**Supplementary Tables**

**Supplementary table 1.** Expression of regional otic vesicle markers in vivo.
**Supplementary table 2.** Genes differentially expressed in each cluster of the otic epithelium subset.
**Supplementary table 3.** Summary of the expression of otic markers in 2D.
**Supplementary table 4.** Raw data and statistics for the expression of otic markers in 3D.
**Supplementary table 5.** Analysis of the directionality of patterning markers in 3D.
